# Single-molecule m^6^A profiling reveals position-dependent mRNA regulation and non-canonical roles for Ythdf2 in early embryogenesis

**DOI:** 10.64898/2026.07.03.736379

**Authors:** Sarah A. Alshawi, Anna Delgado-Tejedor, Srihari Madhavan, Laia Llovera, Rebeca Medina, Cassandra Kontur, Eva Maria Novoa, Jean-Denis Beaudoin

## Abstract

The maternal-to-zygotic transition (MZT) requires coordinated clearance and deadenylation of maternally deposited mRNAs, yet the underlying molecular mechanisms remain poorly understood. N6-methyladenosine (m^6^A) has emerged as a key regulator of maternal mRNA fate, but prior studies have relied on population-averaged short-read methods that cannot resolve modification state, poly(A) tail length, or isoform identity on the same molecule. Here, we employ nanopore direct RNA sequencing on zebrafish embryos across MZT to resolve the interplay between m^6^A deposition, mRNA clearance, and poly(A) tail length dynamics at single-molecule resolution. We find that 78% of expressed maternal genes harbor m^6^A-modified isoforms, significantly exceeding prior bulk estimates. Within-isoform comparisons demonstrate that m^6^A promotes mRNA decay, with CDS m^6^A contributing more to maternal mRNA clearance than 3’-UTR m^6^A. The positional context of m^6^A alone is sufficient to determine the temporal regulation of poly(A) tail lengths. CDS m^6^A constitutively suppresses tail length throughout MZT, while 3′-UTR m^6^A acquires shortening activity only after zygotic genome activation (ZGA). Transcriptomic analysis of *ythdf2* knockout embryos reveals two unrecognized roles. Ythdf2 stabilizes m^6^A-marked maternal transcripts to set stoichiometry at MZT onset, and is also responsible for maintaining global poly(A) tail homeostasis prior to ZGA through an m^6^A-independent mechanism. Together, these findings define the single-molecule logic by which m^6^A modifications shape transcript fate during vertebrate MZT.

## Introduction

The maternal-to-zygotic transition (MZT) is a developmental transition conserved across metazoans that occurs after fertilization. In MZT, developmental control shifts from an exclusive reliance on maternally deposited gene products to the zygotic genome^1–3^. Successful MZT requires both zygotic genome activation (ZGA) and the concurrent remodeling of the maternal transcriptome through regulated changes in mRNA stability and translation^4–7^. Maternal mRNA fate is determined by multiple post-transcriptional mechanisms^3,8^, including *cis*-regulatory elements in 3′ untranslated regions (3′-UTRs)^9^, dynamic RNA-binding proteins^10^, deadenylation^11,12^, and terminal uridylation^13^. Poly(A) tail length is central to this process^11,14^. In early embryos, tail length is tightly coupled to both translational efficiency and mRNA stability, such that changes in adenylation state can rapidly and selectively reprogram the translatome^15,16^ and destabilize targeted transcripts^17^. For example, cytoplasmic polyadenylation of transcripts containing CPE elements in their 3′-UTRs promotes their translation at specific developmental stages^18–20^, whereas transcripts targeted by the microRNA miR-430 undergo deadenylation and decay during MZT in zebrafish^21^. Understanding how maternal transcripts are selectively stabilized, translated, deadenylated, and cleared is therefore essential for defining the regulatory principle governing MZT.

Among the post-transcriptional mechanisms that shape maternal RNA fate, N6-methyladenosine (m^6^A) has emerged as a major regulatory axis. M^6^A is the most abundant internal modification in eukaryotic mRNA^22^ and is deposited co-transcriptionally at DRACH motifs^23,24^ by a writer complex centered on the METTL3-METTL14 methyltransferase heterodimer, with auxiliary components including WTAP, VIRMA, and ZC3H13^25–31^. Methylation is reversible; both FTO and ALKBH5 have been reported to act as demethylases^32,33^. m^6^A is distributed at DRACH motifs unless physically occluded by the exon junction complex (EJC)^34–36^. The EJC blocks METTL3 access within ∼100 nucleotides of splice junctions and, in effect, concentrates m^6^A in long terminal exons, predominantly in the 3’-UTRs and around the stop codon^34^.

Recent studies have demonstrated that the regional placement of m^6^A has functional consequences. m^6^A modifications, including those in the 3’-UTR, are generally read by YTHDF proteins that recruit deadenylase complexes to drive canonical m^6^A-mediated decay^37,38^. CDS-localized m^6^A can additionally engage the CDS-m^6^A decay (CMD) pathway, a distinct, translation-coupled mechanism in which ribosome pausing at modified codons triggers rapid transcript destabilization^39–41^. m^6^A can also stabilize transcripts through recognition by IGF2BP family proteins^42–44^, establishing a bidirectional regulatory logic in which modification state and reader identity together determine transcript fate. Given that MZT requires both targeted clearance and selective stabilization of maternal mRNAs, this positional and reader-dependent complexity makes m^6^A a strong candidate for orchestrating context-dependent transcript fate decisions during this transition.

Several studies in vertebrates have implicated m^6^A as a critical regulator of maternal mRNA fate across the oocyte-to-embryo transition and early embryogenesis. In mice, oocyte-specific depletion of the writer METTL3 impairs oocyte maturation, disrupts maternal mRNA turnover and ZGA, and compromises developmental competence^45^. Loss of the reader YTHDF2 in oocytes causes female infertility by altering maternal transcript dosage and producing oocytes unable to sustain early zygotic development^46^. Notably, loss of the eraser ALKBH5 also causes female infertility, not by preventing clearance, but by failing to remove m^6^A from a subset of transcripts that are then stabilized through m^6^A reader IGF2BP2^47^, illustrating that m^6^A can promote stability during oocyte maturation. In zebrafish, antibody-based m^6^A profiling of early embryos estimated that more than one-third of maternal mRNAs carry m^6^A^48^. Analysis of a maternal-zygotic *ythdf2* (MZ*ythdf2*) mutant led to the proposal that Ythdf2-mediated decay of methylated maternal transcripts is required for timely MZT progression and ZGA^49^. However, a subsequent study using the same MZ*ythdf2* allele found no disruption to global maternal mRNA clearance, m^6^A-dependent decay, ZGA, or gastrulation onset, while demonstrating that m^6^A promoted maternal mRNA deadenylation^49^. The latter study further showed that combined loss of Ythdf2 and Ythdf3 prevents female gonad formation, whereas triple loss of Ythdf1/2/3 is lethal at the larval stage^49^, indicating that Ythdf paralogs are largely functionally redundant and thus capable of compensating for the loss of any single family member^50^. Critically, neither study could determine whether individual maternal mRNAs are fully, partially, or unmethylated, nor could they link m^6^A status directly to poly(A) tail dynamics on the same molecule, leaving the mechanistic basis of m^6^A-dependent fate decisions unresolved at the single-transcript level.

A major limitation of prior studies examining the role of m^6^A during embryogenesis^51–54^ has been their reliance on antibody-based mapping, which enriches m^6^A-containing RNA fragments by immunoprecipitation prior to next-generation sequencing technologies (m^6^A RIP-seq, miCLIP-seq, and related methods)^23,48,55,56^. Even in highly optimized low-input implementations^57,58^, these approaches detect m^6^A indirectly through IP-versus-input enrichment ratios and define modification at the level of broad peak regions, typically spanning hundreds of nucleotides, rather than at individual sites or individual molecules. As a consequence, they provide a population-averaged view of methylation. Thus, antibody-based approaches are typically unable to report modification stoichiometry at single-nucleotide resolution, cannot determine the modification state of individual RNA molecules, and cannot assess whether multiple m^6^A sites on the same transcript are co-modified^59,60^. Critically, because these methods require RNA fragmentation, they cannot link m^6^A status to other molecule-level features, including poly(A) tail length and isoform identity, within the same sequencing read. In this regard, nanopore direct RNA sequencing (DRS) overcomes these limitations by sequencing full-length native RNA molecules without fragmentation or amplification^61^. When combined with m^6^A-aware basecalling algorithms^59,62^, DRS resolves m^6^A at single-nucleotide and single-molecule (read) resolution, enabling accurate quantification of per-site modification stoichiometry across molecules, while simultaneously preserving transcript isoform identity and measuring poly(A) tail length on each individual native RNA molecule^59,63^. These capabilities make DRS uniquely suited to dissect the molecule-level interplay among m^6^A stoichiometry, positional context, isoform identity, and poly(A) tail dynamics that likely underlies context-dependent maternal mRNA fate decisions during MZT.

Here, we apply DRS to a developmental time course in zebrafish embryos to generate the first single-molecule, multi-feature epitranscriptomic atlas of vertebrate MZT. Three findings illustrate the single-molecule and nucleotide resolution gained. First, we find that the maternal transcriptome is far more broadly methylated than the one-third previously estimated by bulk m^6^A profiling, with most expressed genes harboring methylated transcripts. Second, by directly comparing m^6^A+ and m^6^A- reads of the same isoform, we show that m^6^A promotes maternal mRNA decay and poly(A) tail shortening independently of transcript sequence, an inference that bulk across-transcript comparisons cannot make. Third, we resolve a positional logic invisible to bulk methods: CDS m^6^A constitutively suppresses poly(A) tail length from before ZGA, whereas 3’-UTR m^6^A acquires deadenylation activity only after ZGA, indicating that the same modification directs distinct temporal fates depending on its location. Extending our DRS analyses to MZ*ythdf2* mutants helps reconcile the conflicting conclusions of prior genetic studies^48,49^. We find that Ythdf2 is not the primary effector of m^6^A-dependent decay, as global clearance and deadenylation proceed normally for both m^6^A+ and m^6^A- reads. Yet, we uncover two previously unrecognized roles for the reader that were inaccessible to bulk approaches. We find that Ythdf2 maintains m^6^A stoichiometry and global poly(A) tail homeostasis at the onset of MZT. Together, this first per-read analysis of m^6^A across a developmental transition defines the single-molecule logic by which m^6^A shapes maternal mRNA fate during vertebrate MZT.

## Results

### The zebrafish maternal transcriptome is broadly m^6^A-modified with methylated genes exhibiting longer poly(A) tails and higher translation efficiency

To examine how m^6^A is associated with maternal mRNA fate during zebrafish MZT, we performed direct RNA-sequencing by nanopore (DRS) on poly(A)-selected RNA from zebrafish embryos at 2, 4, and 6 hours post-fertilization (hpf), in biological duplicates (**Fig. 1A**). These time points were selected to capture transcriptomic states spanning the onset, peak, and resolution of MZT. In zebrafish, ZGA occurs in two waves: a minor wave beginning around 2 hpf that includes early activation of the miR-430 locus, and a major wave at the mid-blastula transition around 3 hpf^64,65^. By 6 hpf, zygotic transcription is well established and maternal mRNA clearance is underway through deadenylation, uridylation, and miR-430-mediated decay^13,21^ (**Fig. 1A**). DRS provides molecule-resolved measurements of m^6^A status, poly(A) tail length, and isoform identity for each sequenced mRNA, with the long-read nature of each sequencing read enabling unambiguous assignment to specific isoforms. Replicates showed strong concordance, with per-isoform read counts and mean poly(A) tail lengths showing Pearson correlations of 0.94-0.99 and 0.90-0.99, respectively, across time points (**Supp. Fig. 1A,B**). DRS poly(A) tail lengths were in agreement with PAL-seq^15^ (**Supp. Fig. 1C**), an orthogonal technique for tail length measurement. m^6^A sites were identified using an m^6^A-aware basecalling workflow validated for per-read, single-nucleotide m^6^A prediction from native RNA^59^ (see Methods).

**Figure 1.**
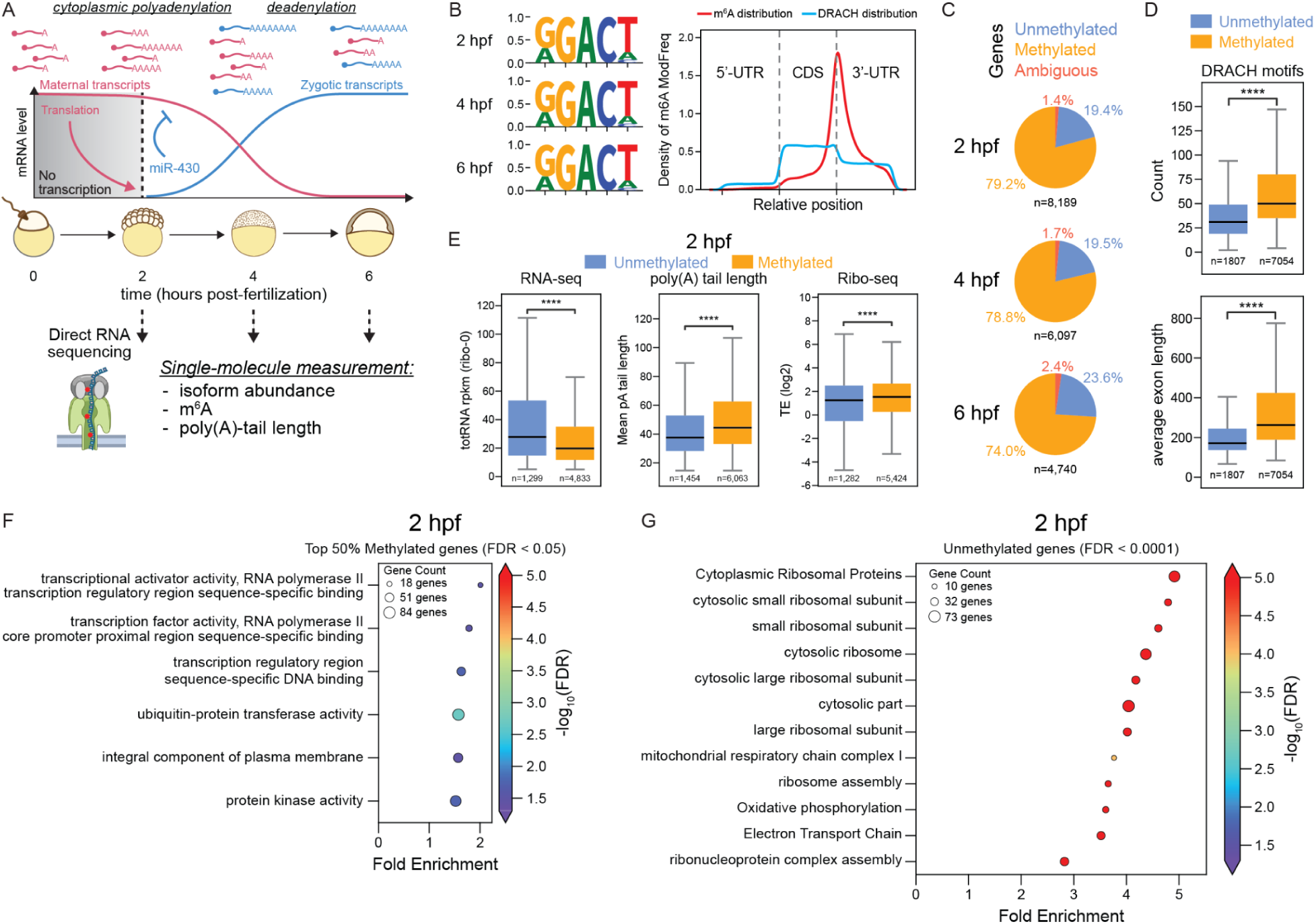
Single-molecule m^6^A profiling of the maternal transcriptome reveals a large population of methylated mRNAs with distinct expression, poly(A) tail, and translation properties. **(A)** Schematic of transcript dynamics during zebrafish MZT, with the DRS time course and single-molecule measurements indicated. **(B)** Sequence motif enriched at m^6^A sites at 2, 4, and 6 hpf (left), and metagene distribution profiles of m^6^A modifications (red) and DRACH motifs (blue) across the 5’-UTR, CDS, and 3’-UTR (right). (**C**) Percentage of genes classified as methylated (≥4 m^6^A+ reads), unmethylated (0 m^6^A+ reads), or ambiguous (1-3 m^6^A+ reads) at each timepoint. (**D**) Average number of DRACH motifs per isoform (upper) and average exon length (lower) for unmethylated and methylated isoforms at 2 hpf (Mann-Whitney tests, **** P<0.0001). (**E**) Expression level (RNA-seq), poly(A) tail length, and translation efficiency (Ribo-seq) of unmethylated and methylated isoforms at 2 hpf (Mann-Whitney tests, **** P<0.0001). (**F**) Gene ontology fold enrichment for the top 50% most heavily methylated gene population at 2 hpf (FDR < 0.05) (**G**) Gene ontology fold enrichment for unmethylated gene population at 2 hpf (FDR < 0.0001).

We first asked whether the m^6^A calls reproduced canonical features of transcriptome-wide m^6^A deposition. Across the time course, methylated sites were enriched in a DRACH-like motif and accumulated preferentially in the CDS and 3′ UTR, with a pronounced peak near the stop codon (**Fig. 1B**). This topology is consistent with m^6^A distributions previously described using DRS in human cell lines^59,66^, and with antibody-based profiling in zebrafish embryos^48^. Notably, m^6^A enrichment near the stop codon exceeded that expected from DRACH motif occurrence alone (**Fig. 1B**), consistent with EJC-mediated exclusion of methylation from splice junction-proximal regions concentrating m^6^A toward long terminal exons^34–36^.

Because prior studies of m^6^A during early development used bulk antibody-based methods that resolve methylation at the population rather than single-molecule level^51–54^, we summarized our single-molecule calls at the gene level to enable direct comparison. Using a conservative threshold, defining genes with ≥4 m^6^A-containing reads as methylated, 0 reads as unmethylated, and 1-3 reads as ambiguous, we found that more than 78.4% of expressed maternal genes harbored methylated mRNAs at 2 hpf, with the proportion declining modestly but consistently across MZT to 74% at 6 hpf (**Fig. 1C**). This prevalence is substantially higher than the one-third estimate from earlier bulk m^6^A-seq and m^6^A-CLIP-seq profiling of zebrafish embryos^48^, likely reflecting the improved sensitivity of DRS in detecting low-stoichiometry sites and avoiding the sequence and length biases inherent to antibody-based enrichment. Isoforms expressed from unmethylated genes appear structurally predisposed against methylation, characterized by shorter mRNA length, fewer DRACH motifs, particularly outside the 100 nucleotide exclusion zone flanking splice junctions, and shorter average exon lengths (**Fig. 1D**, **Supp. Fig. 1D,E**), features collectively predicted to reduce METTL3 accessibility and m^6^A deposition. The decline in the fraction of methylated genes across MZT is consistent with progressive clearance of m^6^A-marked mRNAs, a dynamic we examine at single-molecule resolution in subsequent analyses.

Despite representing the minority class, unmethylated isoforms were on average more highly expressed than methylated isoforms pre-ZGA, both in independently generated short-read RNA-seq data^9^ and in the DRS dataset (**Fig. 1E**, left panel**; Supp. Fig. 1F**). This challenges the possibility that the unmethylated classification simply reflects lower sequencing coverage relative to methylated isoforms. By contrast, methylated isoforms exhibited longer poly(A) tails than unmethylated isoforms in our dataset and higher translation efficiency in a previously published ribosome profiling dataset^67^ (**Fig. 1E**, middle and right panels). This is consistent with tail-length-dependent translational regulation in the pre-ZGA embryo, where limiting cytoplasmic PABP availability causes longer-tailed mRNAs to more effectively compete for ribosome recruitment^16^. m^6^A-marked maternal isoforms thus appear to be preferentially maintained in a translationally competent state at the onset of MZT. To ask whether these two populations also differ in the biological processes they encode, we performed gene ontology analysis at 2 hpf. Genes with isoforms in the top 50% of m^6^A stoichiometry were enriched for transcriptional regulatory functions, including RNA polymerase II transcription factor activity, transcription regulatory region binding, ubiquitin-protein transferase activity, and protein kinase activity (**Fig. 1F**). In contrast, unmethylated genes were enriched for cytosolic ribosome structure and assembly, oxidative phosphorylation, mitochondrial respiratory chain complex I, and electron transport chain components (**Fig. 1G**). This functional dichotomy suggests that m^6^A preferentially marks transcripts encoding developmental regulators and signaling proteins, categories subject to tight spatiotemporal control during MZT, while core housekeeping machinery supporting translation and energy metabolism is largely excluded from methylation.

### DRS captures isoform-level decay dynamics and deadenylation patterns across MZT

Timely clearance of maternal mRNAs through both maternally- and zygotically-encoded decay mechanisms is essential for MZT progression^1–6^. To validate that our DRS dataset faithfully captures established decay biology, we classified expressed mRNAs into the four categories defined by Vejnar *et al.*^9^: maternally-encoded decay (M-decay), miR-430-mediated decay (430-decay), zygotically-encoded decay (Z-decay), and an “other” category comprising mRNAs that do not undergo significant decay across MZT or are primarily of zygotic origin. Briefly, M-decay mRNAs are degraded independently of ZGA, miR-430 targets are stabilized by locked nucleic acid (LNA)-mediated miR-430 inhibition, and Z-decay mRNAs are stabilized by ZGA inhibition but insensitive to miR-430 blockade^9^. All three decay populations showed progressive downregulation across the time course, with the most pronounced changes occurring between 4 and 6 hpf (**Fig. 2A and 2B**). This is consistent with previously reported clearance timing^9^. Conversely, mRNAs increasing in abundance across MZT were largely confined to the “other” category (**Fig. 2B**), as expected for mRNAs of zygotic origin whose accumulation reflects ZGA rather than maternal transcript dynamics.

**Figure 2.**
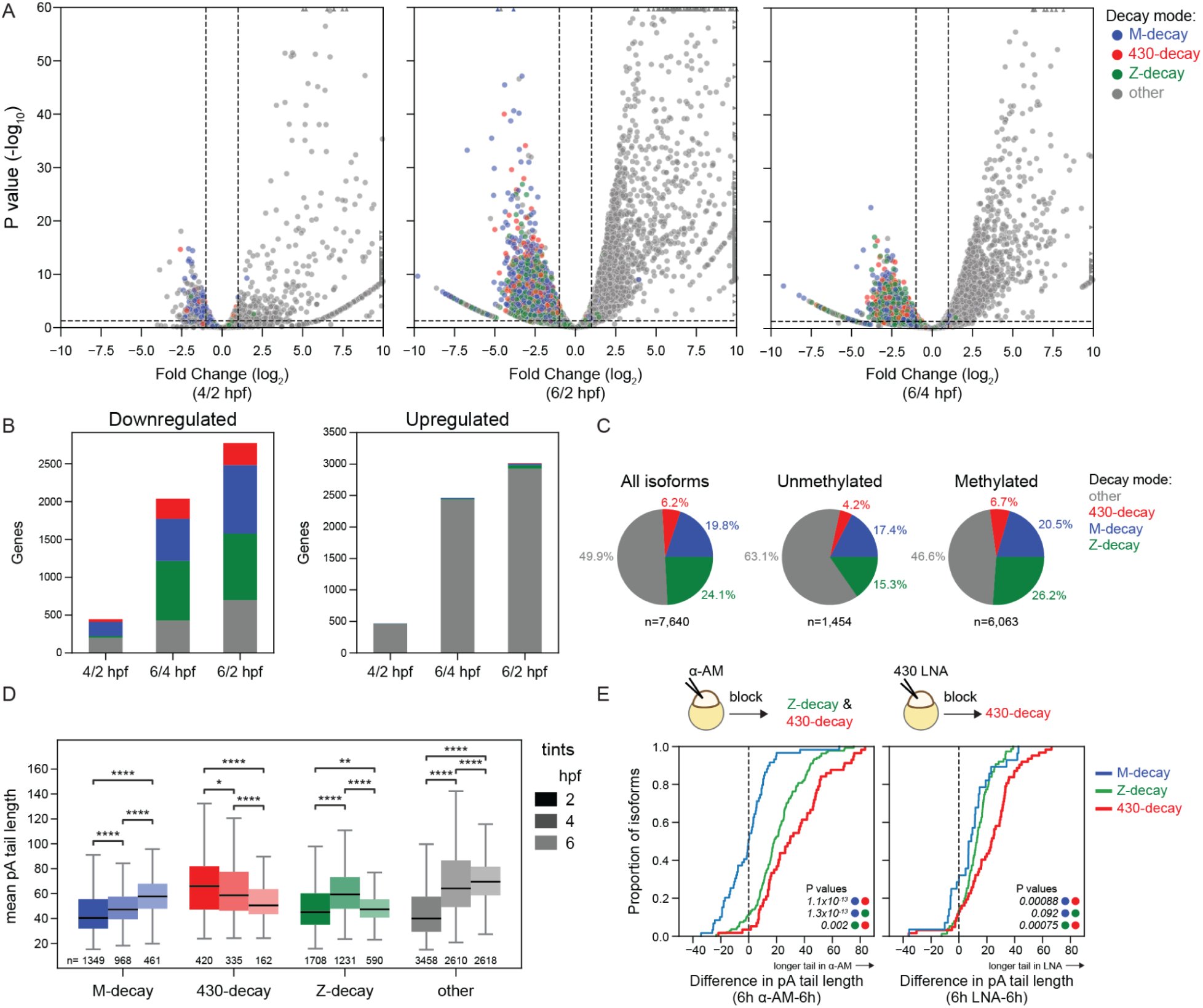
DRS captures decay-mode-specific clearance and deadenylation, with methylated genes enriched in zygotically-controlled decay pathways. (**A**) Volcano plots of differential expression between consecutive time points (4/2, 6/2, and 6/4 hpf), showing log_2_ fold change versus –log_10_ adjusted *P*-value (DESeq2). Genes are colored by decay mode (M-decay, blue; 430-decay, red; Z-decay, green; other, gray). (**B**) Number of significantly downregulated (left) and upregulated (right) genes at each timepoint comparison, colored by decay mode. (**C**) Proportion of isoforms assigned to each decay mode among all, unmethylated, and methylated isoforms. (**D**) Mean poly(A) tail length of isoforms in each decay mode (color) at 2, 4, and 6 hpf (tint). Black lines indicate medians, boxes span the firsts (Q1) to third (Q3) quartiles, and whiskers extend to Q1 -1.5×IQR and Q3 + 1.5×IQR. N values are indicated below each box. Statistical significance was assessed by Mann-Whitney tests with Benjamini-Hochberg correction (* P<0.05, ** P<0.01, *** P<0.001, **** P<0.0001). (**E**) Cumulative distributions of the change in poly(A) tail length following α-amanitin treatment (left) or miR-430 LNA inhibition (right) at 6 hpf. P-values were calculated by Mann-Whitney U tests between the indicated categories.

Having confirmed that our dataset captures previously reported global decay trends, we next examined how m^6^A-methylated genes are distributed across decay modes. Consistent with prior evidence that m^6^A and miR-430 act through independent but partially overlapping mechanisms to promote maternal mRNA clearance^49^, methylated genes were enriched across 430-, Z- and M-decay modes relative to unmethylated genes (**Fig. 2C**, **Supp. Fig. 2A,B**). This enrichment was most pronounced in the 430-decay (*P*=9.56x10⁻^4^) and Z-decay (*P*=4.58x10⁻^17^) populations (**Supp. Fig. 2A,B**), suggesting that mRNAs subject to zygotically-encoded clearance mechanisms are preferentially m^6^A-marked, a pattern that positions m^6^A as a co-occurring feature of mRNAs destined for post-ZGA clearance.

We next examined how poly(A) tail lengths in each decay population change across the MZT time course. miR-430 targets carried the longest tails at 2 hpf and underwent progressive poly(A) tail shortening through 6 hpf (**Fig. 2D**, **Supp. Fig. 2C**), directly reflecting the progression of miR-430-mediated deadenylation as ZGA proceeds and miR-430 levels rise. M-decay mRNAs showed a progressive increase in tail length across the time course, while Z-decay mRNAs showed an increase in tail length between 2 and 4 hpf followed by pronounced shortening between 4 and 6 hpf, coinciding with the major wave of zygotic factor-mediated clearance (**Fig. 2D**, **Supp. Fig. 2C**). These poly(A) tail length dynamics are consistent with those independently measured when validating Nano3P-seq, an orthogonal cDNA-based nanopore sequencing approach that captures transcripts regardless of polyadenylation status^68^, providing additional validation of our DRS-based poly(A) tail measurements. Notably, 430-, M-, and Z-decay mRNAs maintained shorter tails than the “other” category post-ZGA, reflecting their ongoing deadenylation relative to the characteristically longer tails of the predominantly zygotically derived “other” mRNAs (**Supp. Fig. 2C**). Together, these temporal dynamics establish that our DRS time course captures decay-mode-specific deadenylation trajectories at isoform resolution.

We then confirmed that the progressive poly(A) tail shortening observed in miR-430 targets specifically reflects miR-430-dependent deadenylation. Treatment with α-amanitin, which blocks RNA polymerase II preventing both ZGA and zygotic miR-430 production^9,69^, selectively increased poly(A) tail lengths in the 430- and Z-decay populations but not in M-decay mRNAs at 6 hpf (**Fig. 2E**, left panel). Conversely, LNA-mediated inhibition of miR-430 alone^70^ selectively increased tail lengths in the 430-decay population without affecting Z-decay mRNAs (**Fig. 2E**, right panel). These results establish that DRS-based poly(A) measurements faithfully capture deadenylation in a decay-mode-specific manner at isoform resolution, providing a validated framework for interrogating the role of m^6^A in regulating poly(A) tail dynamics and transcript fate during MZT.

### m^6^A promotes transcript decay and deadenylation in an mRNA sequence-independent manner

Given that DRS allows for single-molecule resolution unlike previously used methods, we then characterized transcript-level m^6^A patterns during MZT. We quantified the fraction of reads carrying at least one m^6^A modification (m^6^A+ reads) for each methylated isoform across the time course. On average, 35.2% of reads per methylated isoform were m^6^A+ at 2 hpf, declining modestly to 32.3% by 6 hpf (**Fig. 3A**). This average across methylated isoforms, however, masks substantial heterogeneity in modification stoichiometry across individual isoforms. While many methylated isoforms carry m^6^A on only a small fraction of their reads, others are modified on the large majority, with the m^6^A+ fraction reaching >94% and 126 isoforms exceeding 75% m^6^A+ reads at 2 hpf (**Fig. 3B**, **Supp. Fig. 3A,B** and **Supp. Dataset 1**). Additionally, our data revealed that most m^6^A+ reads contain only one m^6^A modification (**Supp. Fig. 3C**). Supporting the validity of these measurements, isoforms previously identified as methylated by antibody-based approaches showed higher m^6^A+ read fractions than those previously categorized as unmethylated^48,49^ (**Supp. Fig. 3D**).

**Figure 3.**
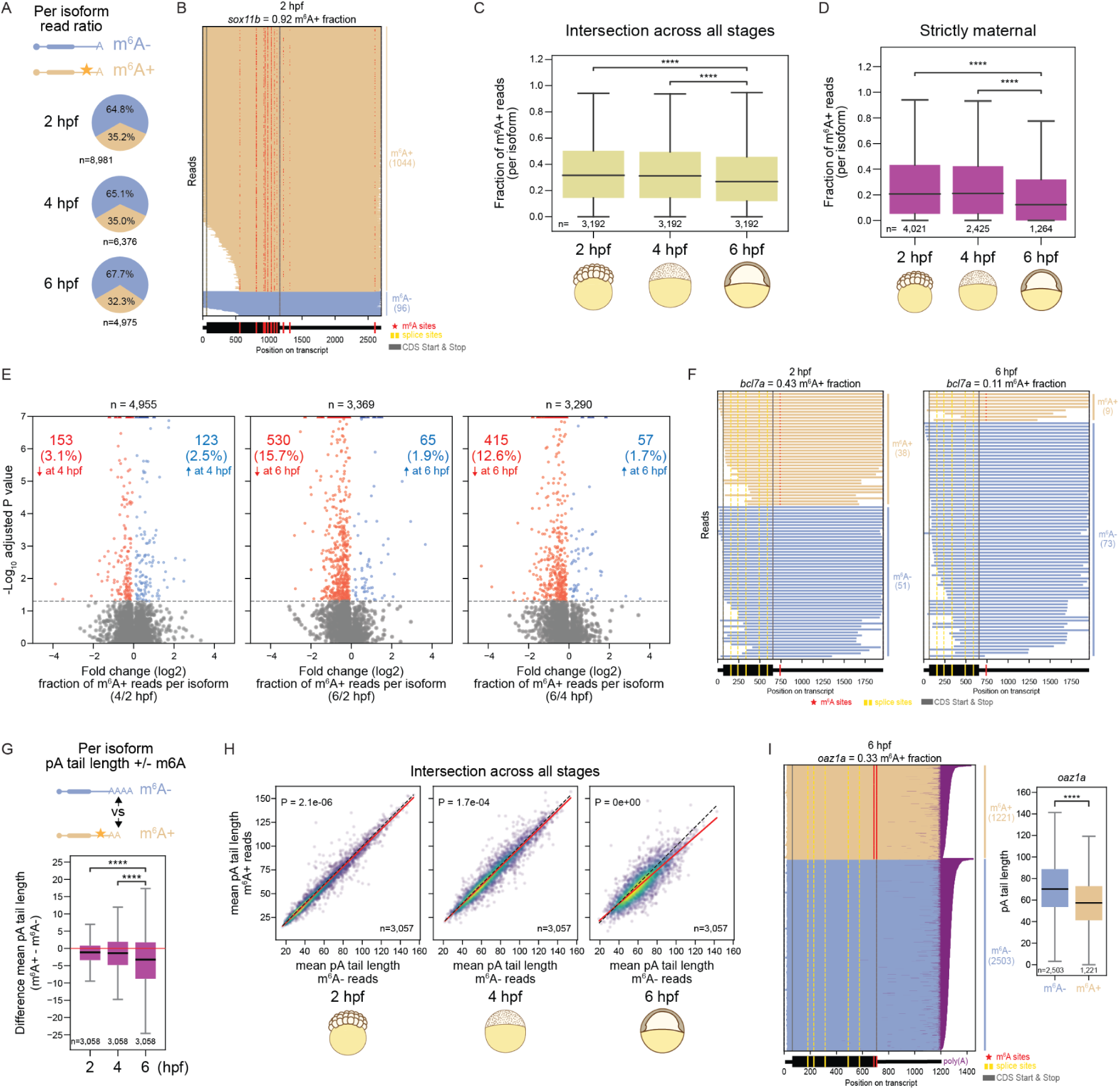
Within-isoform single-molecule analysis links m^6^A status to mRNA decay and deadenylation during MZT. (**A**) Pie charts showing the average fraction of reads per methylated isoform that carry m^6^A (m^6^A+) or lack m^6^A (m^6^A-) at 2, 4, and 6 hpf. (**B**) Read-level view of a representative isoform *sox11b* (ENSDART00000141068) with a high m^6^A+ fraction at 2 hpf. Each row is one read (m^6^A+, brown; m^6^A-, blue), with m6A sites (red), splice sites (yellow), and CDS start/stop (gray). (**C**) Fraction of m^6^A+ reads per isoform across isoforms detected at all three time points (2, 4, and 6 hpf). Only uniquely mapping reads were included, and isoforms were required to have at least 25 reads per replicate. Mann-Whitney U tests with Bonferroni correction were used to assess significance between time points (**** P<0.0001). (**D**) Fraction of m^6^A+ reads per isoform for strictly maternal isoforms with no detectable zygotic contribution^71^. Mann-Whitney tests with Bonferroni correction were used to assess significance between time points (**** P<0.0001). Only uniquely mapping reads were included, and isoforms were required to have at least 25 reads per replicate. (**E**) Volcano plots of the change in m^6^A+ read fraction per isoform between time points (4/2, 6/2, 6/4 hpf). The horizontal dashed line marks adjusted P = 0.05. Isoforms with a significant decrease in m^6^A+ fraction are shown in red (count and percentage, upper left), and those with a significant increase in blue (upper right). P-values were calculated by chi-square test with Benjamini-Hochberg correction; points beyond the axis limits are shown as triangles. (**F**) Read-level views at 2 and 6 hpf of a representative isoform *bcl7a* (ENSDART00000190477) that decreases in m^6^A+ fraction across the time course. (**G**) Difference in mean poly(A) tail length for m^6^A+ and m^6^A-reads of isoforms detected at all three time points. Significance between timepoints calculated by unpaired t-test. (**** P<0.0001) (**H**) Per-isoform scatter plots comparing the mean poly(A) tail length of m^6^A+ reads versus m^6^A- reads at 2, 4, and 6 hpf, for isoforms detected at all three time points. Each point represents one isoform and color indicates point density. The red line is the regression fit and the dashed line is the identity (y = x) line. P-values are shown for the difference between the regression and identity lines (see Methods for calculation details). (**I**) Read-level view of a representative isoform *oaz1a* (ENSDART00000105532) at 6 hpf with poly(A) tails shown (purple), alongside box plots comparing mean poly(A) tail length of m^6^A+ versus m^6^A- reads. P-value calculated by Mann-Whitney U test (**** P<0.0001).

Prior work in zebrafish has implicated m^6^A in promoting maternal transcript decay during MZT^48,49^. We reasoned that if m^6^A promotes decay, the fraction of m^6^A+ reads per isoform should decrease over MZT as m^6^A-marked molecules are preferentially cleared. Critically, by comparing m^6^A+ and m^6^A- reads within the same isoform rather than across genes, we control for all sequence-intrinsic features, including 3′-UTR composition, codon usage, and miRNA target sites, that independently influence mRNA stability. This within-isoform comparison therefore isolates the contribution of m^6^A modification status to transcript fate. Consistent with preferential decay of m^6^A-containing molecules, the fraction of m^6^A+ reads per isoform decreased significantly at 6 hpf (**Fig. 3C**). This trend was also observed in strictly maternal genes without detectable zygotic contribution^71^, indicating that the declining m^6^A+ fraction reflects preferential decay of methylated molecules rather than dilution by newly synthesized unmethylated zygotic transcripts (**Fig. 3D, Supp. Fig. 4A**). At the isoform level, 530 (15.7%) and 415 (12.6%) isoforms showed a statistically significant decrease in their m^6^A+ read fraction between 2-6 hpf and 4-6 hpf, respectively (**Fig. 3E,F**). Isoforms with a statistically significant decline in m^6^A+ read fraction between 4 and 6 hpf were enriched in genes localizing to the nucleolar compartment (**Supp. Fig. 4B**), suggesting that m^6^A-dependent clearance may be particularly active for this functional class of maternal mRNAs, though the mechanism of this enrichment remains to be determined.

**Figure 4.**
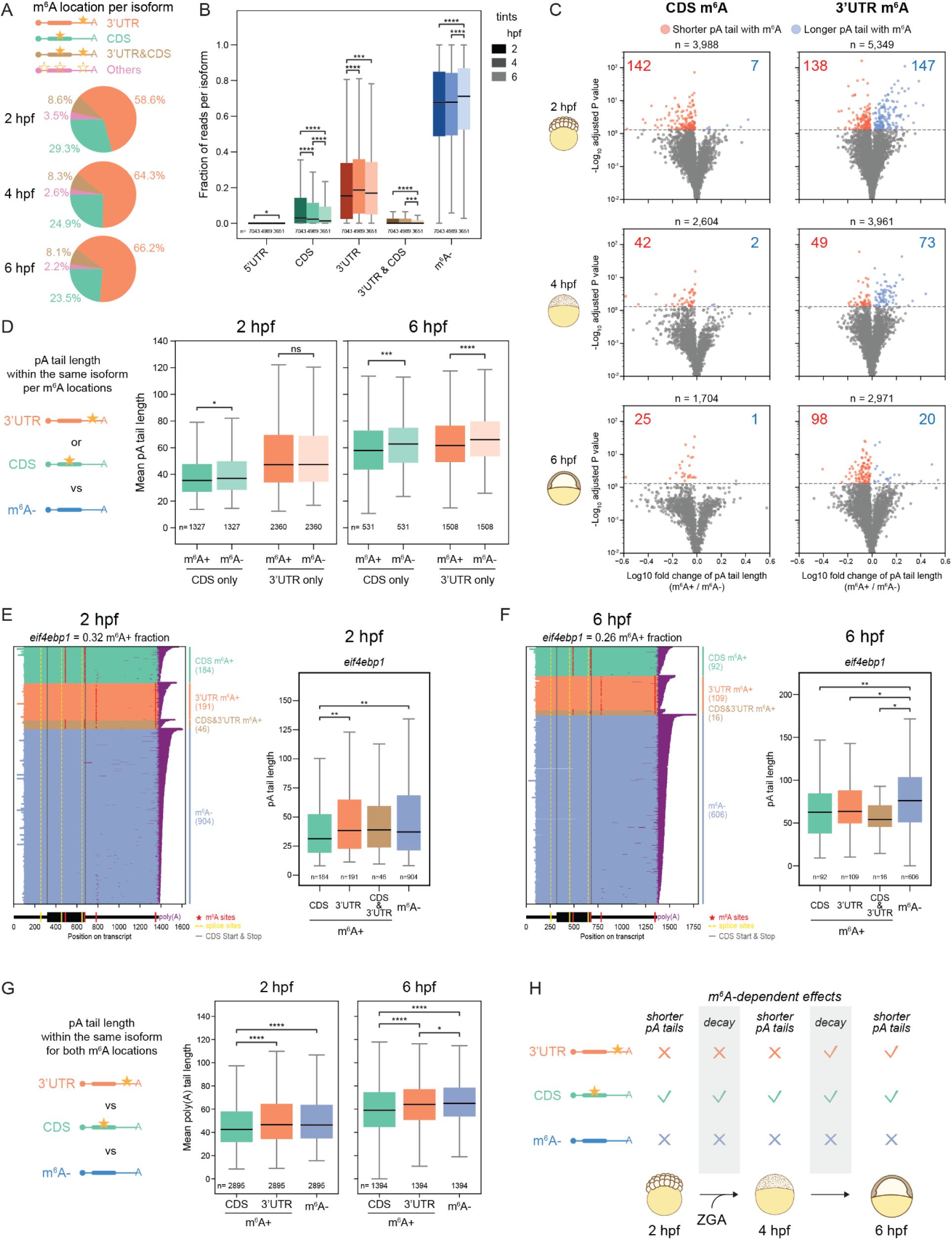
CDS and 3’-UTR m^6^A exert temporally distinct effects on mRNA decay and poly(A) tail length across MZT. (**A**) Pie charts showing the average fraction of m^6^A+ reads carrying m^6^A in each transcript region (3′-UTR, CDS, both 3′-UTR and CDS, or other) per methylated isoform at 2, 4, and 6 hpf. (**B**) Fraction of reads per methylated isoform carrying m^6^A in each region (5′-UTR, CDS, 3′-UTR, or both CDS and 3′-UTR) or lacking m^6^A (m^6^A-), across all methylated isoforms at each time point. Box tint indicates time point (2, 4, and 6 hpf). n indicates the number of isoforms per category per time point. Mann-Whitney tests with Bonferroni correction were used to assess significance between time points (* P<0.05, *** P<0.001, **** P<0.0001). (**C**) Volcano plots comparing poly(A) tail length between regional m^6^A+ reads and the corresponding m^6^A- reads of the same isoform, for CDS m^6^A (left) and 3′-UTR m^6^A (right) at 2, 4, and 6 hpf. Isoforms with significantly shorter poly(A) tails in m^6^A+ reads are shown in red (count, upper left) and those with significantly longer tails in blue (count, upper right). P-values were calculated by unpaired t-test between m^6^A+ and m^6^A- reads and adjusted by the Benjamini-Hochberg procedure. The horizontal dashed line marks adjusted P = 0.05, and points beyond the axis limits are shown as triangles. (**D**) Mean poly(A) tail length of m^6^A+ and corresponding m^6^A- reads for isoforms with m^6^A in the CDS only or 3′-UTR only, at 2 hpf (left) and 6 hpf (right). Only isoforms carrying CDS m^6^A+ reads without any 3′-UTR m^6^A+ reads (or vice versa) were included. n indicates the number of isoforms per category. Mann-Whitney U tests were used to assess significance (ns, not significant; * P<0.05, ** P<0.01). (**E**) Read-level view of a representative isoform *eif4ebp1* (ENSDART00000064032) at 2 hpf, with each row representing a read colored by m^6^A location (CDS m^6^A+, green; 3’-UTR m^6^A+, orange; CDS and 3’UTR m^6^A+, tan; m^6^A-, blue). m^6^A sites (red), splice sites (yellow), CDS start and stop (gray), and poly(A) tails (purple) are indicated, with read counts per category on the right (left panel). Box plot of per-read poly(A) tail length by m^6^A location for *eif4ebp1* (right panel). (**F**) Same as (E) for *eif4ebp1* at 6 hpf (left). Boxplot of *eif4ebp1* per read poly(A) tail length by m^6^A location at 6hpf (right). Statistical significance was assessed by Mann-Whitney U test (* P<0.05, ** P<0.01). (**G**) Mean poly(A) tail length within isoforms carrying m^6^A in both the CDS and 3′-UTR, comparing reads methylated in the CDS only, the 3′-UTR only, or lacking m^6^A (m^6^A-), at 2 hpf (left) and 6 hpf (right). *n* values are indicated below each box. Statistical significance was assessed by Mann-Whitney U test (* P<0.05, **** P<0.0001). (**H**) Schematic summarizing the temporally distinct, positional effects of m^6^A on mRNA fate across MZT. CDS m^6^A+ transcripts show shorter poly(A) tails at all three time points and undergo decay between 2-4 hpf and 4-6 hpf. Meanwhile, 3’-UTR m^6^A+ transcripts acquire shorter poly(A) tails only after ZGA, from 4 to 6 hpf.

We next asked whether m^6^A status is associated with poly(A) tail length differences within the same isoform. Prior assessments of m^6^A effects on poly(A) tail length during zebrafish MZT have relied on integrating independent bulk m^6^A and poly(A) tail datasets or on artificial reporter systems^49^, neither of which permits direct within-molecule comparisons on endogenous transcripts. Comparing mean poly(A) tail lengths of m^6^A+ and m^6^A- reads within the same isoforms, we found similar tail lengths at 2 or 4 hpf (**Fig. 3G,H**). By 6 hpf, however, m^6^A+ reads exhibited shorter poly(A) tails than their m^6^A- counterparts from the same isoform (**Fig. 3G,H,I**; **Supp. Fig. 4C-E**). This tail length difference was significantly larger at 6 hpf compared to previous time points (**Fig. 3G),** indicating that preferential deadenylation of m^6^A-containing molecules coincides with the onset of the major wave of maternal mRNA clearance. Together, these within-isoform comparisons directly associate m^6^A modification status with both decay and deadenylation during MZT, independent of confounding sequence differences between transcripts.

### m^6^A location dictates the temporal dynamics of its effect on mRNA decay and poly(A) tail length

Recent work in cultured cells has shown that m^6^A location within a transcript is a key determinant of its functional outcome, with CDS-localized m^6^A promoting translation-coupled decay through ribosome pausing^39–41^, while 3′-UTR m^6^A primarily promotes canonical YTHDF-mediated deadenylation^37,38^. Whether these positional distinctions operate during vertebrate early development is currently unknown. We characterized the distribution of m^6^A across transcript regions throughout MZT. Across the time course, the average distribution of m^6^A per isoform was predominantly in the 3′-UTR and CDS, with a subset of isoforms carrying modifications in both regions simultaneously (**Fig. 4A**). To examine how modification stoichiometry evolves by region across the time course, we quantified the fraction of m^6^A+ reads in each regional category for all expressed isoforms at each time point. As expected from the overall decline in m^6^A+ read fraction described above (**Fig. 3C-E** and **Supp. Fig. 5A**), the global fraction of m^6^A- reads increased at 6 hpf relative to earlier time points (**Fig. 4B**). Strikingly, this increase was driven predominantly by a consistent decline in CDS m^6^A+ reads across all three time points, while the fraction of 3′-UTR m^6^A+ reads initially increased between 2 and 4 hpf before decreasing between 4 and 6 hpf (**Fig. 4B**). This pattern was preserved when the analysis was restricted to strictly maternal mRNAs (**Supp. Fig. 5A**), confirming that the predominance of CDS m^6^A loss reflects differential positional decay dynamics rather than dilution by newly synthesized zygotic transcripts. Together, these results implicate CDS m^6^A as the primary positional contributor to m^6^A-dependent maternal mRNA decay during MZT, consistent with a model in which rising global translation^2,3,7^ progressively accelerates translation-coupled degradation of CDS-methylated transcripts.

**Figure 5.**
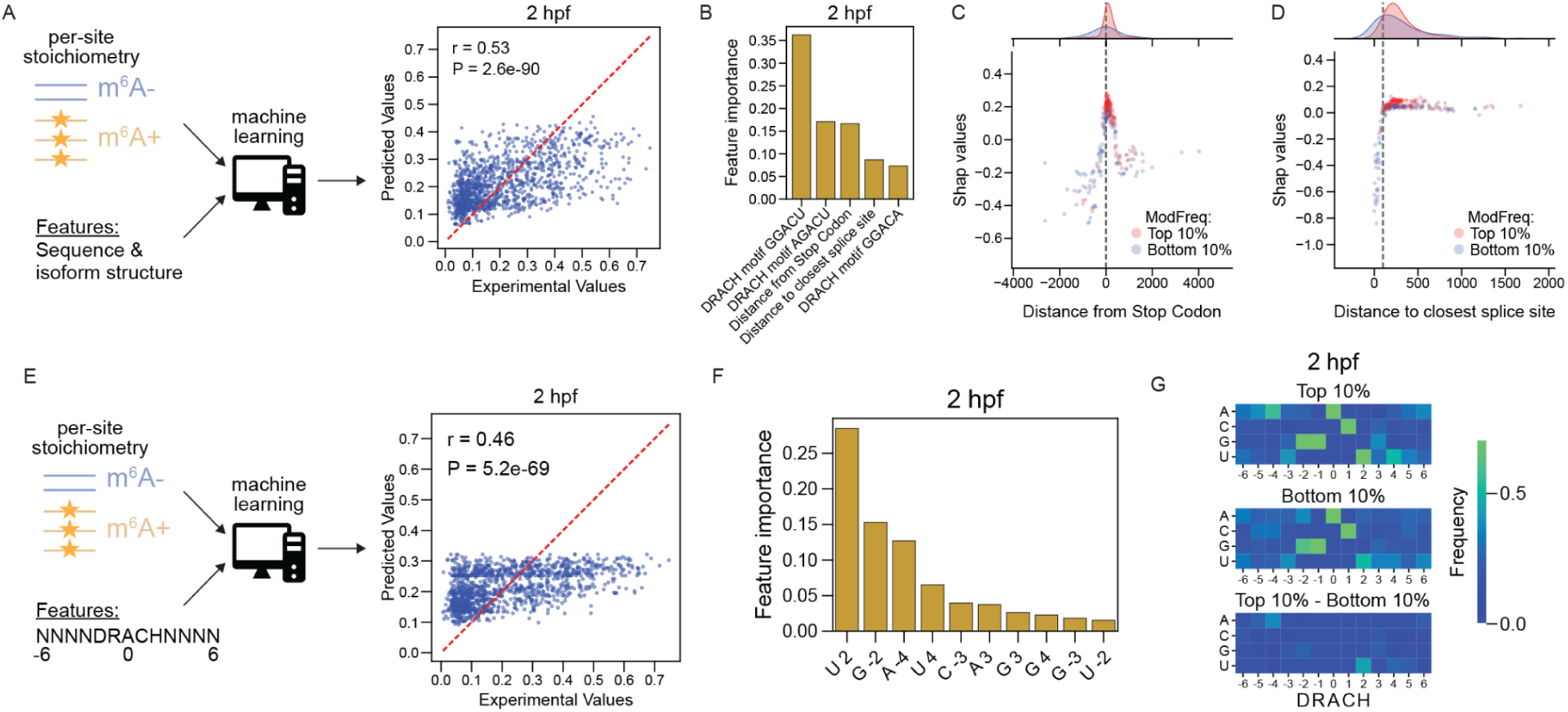
Transcript architecture and local sequence context jointly predict m^6^A stoichiometry across MZT. (**A**) Schematic of the machine learning model trained to predict per-site m^6^A stoichiometry from sequence and isoform structure features, with the predicted versus experimental stoichiometry values at 2 hpf shown (right). The red dashed line indicates the identity (y = x) line. The correlation coefficient (r) and P-value are derived from Spearman correlation. (**B**) Top features ranked by importance in the structure-and-sequence model at 2 hpf. (**C**) SHAP values as a function of distance from the stop codon for sites in the top 10% (red) and bottom 10% (blue) of stoichiometry, with density distributions shown above. (**D**) SHAP values as a function of distance to the nearest splice site for the same top and bottom 10% stoichiometry groups. (**E**) Schematic of the second model, trained to predict per-site stoichiometry using only the DRACH motif and flanking nucleotides, with predicted versus experimental values at 2 hpf shown (right). (**F**) Top features ranked by importance in the DRACH-sequence model at 2 hpf, labeled by nucleotide and position relative to the methylated A. (**G**) Nucleotide frequency at each position flanking the m^6^A site for top 10% (upper) and bottom 10% (middle) stoichiometry sites, and the frequency difference between the two groups (lower). Color indicates nucleotide frequency.

To directly assess the impact of m^6^A location on poly(A) tail length at the single-molecule level, we compared mean poly(A) tail lengths of m^6^A+ and m^6^A- reads within the same isoform, stratified by the region of m^6^A deposition. At the individual isoform level, CDS m^6^A was consistently associated with shorter poly(A) tail lengths, with 142 isoforms showing significantly shorter tails in m^6^A+ reads at 2 hpf compared to only 7 with longer tails, a trend that persisted throughout MZT (**Fig. 4C**). Among isoforms with 3′-UTR m^6^A, the effect on poly(A) tail length was mixed at 2 hpf, with 138 and 147 isoforms showing significantly shorter and longer tails in m^6^A+ reads relative to m^6^A- reads from the same isoforms, respectively (**Fig. 4C**), suggesting that 3′-UTR m^6^A does not uniformly shorten or lengthen poly(A) tails at this early stage. By 6 hpf, the majority of isoforms with 3′-UTR m^6^A-mediated tail length differences showed shorter poly(A) tails in m^6^A+ reads (98 of 118 isoforms), and were enriched for genes involved in RNA splicing and RNA binding (**Supp. Fig. 5B**).

These positional and temporal effects were confirmed at the population-level. At 2 hpf, m^6^A+ reads of isoforms with m^6^A sites exclusively in the CDS (CDS-only) exhibited significantly shorter poly(A) tails than their m^6^A- counterparts, whereas isoforms with exclusively 3′-UTR m^6^A sites (3’-UTR-only) showed no significant poly(A) tail difference (**Fig. 4D**). At 6 hpf, the association of CDS-only m^6^A with shorter poly(A) tails was maintained. Likewise, 3′-UTR-only m^6^A became associated with significantly shorter poly(A) tails relative to respective m^6^A- reads at that timepoint (**Fig. 4D**). The same pattern held within individual isoforms. For the *eif4ebp1* isoform, CDS m^6^A+ reads had shorter tails at 2 and 6 hpf (**Fig. 4E,F**), whereas reads methylated in the CDS, the 3’UTR, or both had shorter tails only at 6 hpf (**Fig. 4F**). Extending this analysis to all isoforms carrying m^6^A sites in both regions, reads methylated exclusively in the CDS had shorter poly(A) tails than m^6^A- reads at both time points, while 3’-UTR exclusive m^6^A+ reads showed no significant difference at 2 hpf but significantly shorter tails at 6 hpf (**Fig. 4G**). These results indicate that CDS m^6^A suppresses poly(A) tail length throughout MZT, consistent with its proposed role in translation-coupled decay^41^, while the poly(A) tail-shortening activity of 3′-UTR m^6^A is developmentally acquired, emerging after ZGA and the onset of zygotic effector activity. Altogether, these findings suggest that the position of m^6^A is sufficient to determine the temporal regulation of mRNA decay and poly(A) tail length during MZT, directing identical mRNA sequences to distinct temporal fates in early embryogenesis (**Fig. 4H**).

### DRACH motif, Stop Codon, and Splice site proximity are Key Predictors of m^6^A Stoichiometry across MZT

Given the developmentally regulated roles of m^6^A placement in transcript decay and deadenylation, we next asked what sequence and structural features determine m^6^A stoichiometry across the maternal transcriptome during MZT. To address this, we trained machine learning models to predict per-site stoichiometry from sequence and isoform structure features (see Methods) (**Fig. 5A** and **Supp. Fig. 6**). The model explained a substantial fraction of stoichiometry variation (r = 0.53), with DRACH motif identity, stop codon proximity, and distance to the nearest splice junction emerging as the most important predictors across all time points (**Fig. 5B** and **Supp. Fig. 6**). High-stoichiometry sites were enriched near the stop codon, while low-stoichiometry clustered within ∼100 nucleotides of splice junctions (**Fig. 5C,D** and **Supp. Fig. 6**). This pattern directly supports the EJC-mediated default-methylation model, in which splice junction-proximal sites are occluded from METTL3 while stop codon-proximal DRACH motifs in long terminal exons are most efficiently modified^34–36^. A second machine learning model restricted to the DRACH motifs and the ±4 flanking nucleotides retained substantial predictive power (r = 0.46), identifying flanking positions -4 and +4 as contributors to stoichiometry beyond the minimal motif (**Fig. 5F,G** and **Supp. Fig. 7**). This suggests that METTL3 recognition extends beyond the five-nucleotide DRACH consensus. Together, these analyses show that m^6^A stoichiometry is jointly encoded by transcript architecture and local sequence context, with both determinants stable across MZT.

**Figure 6.**
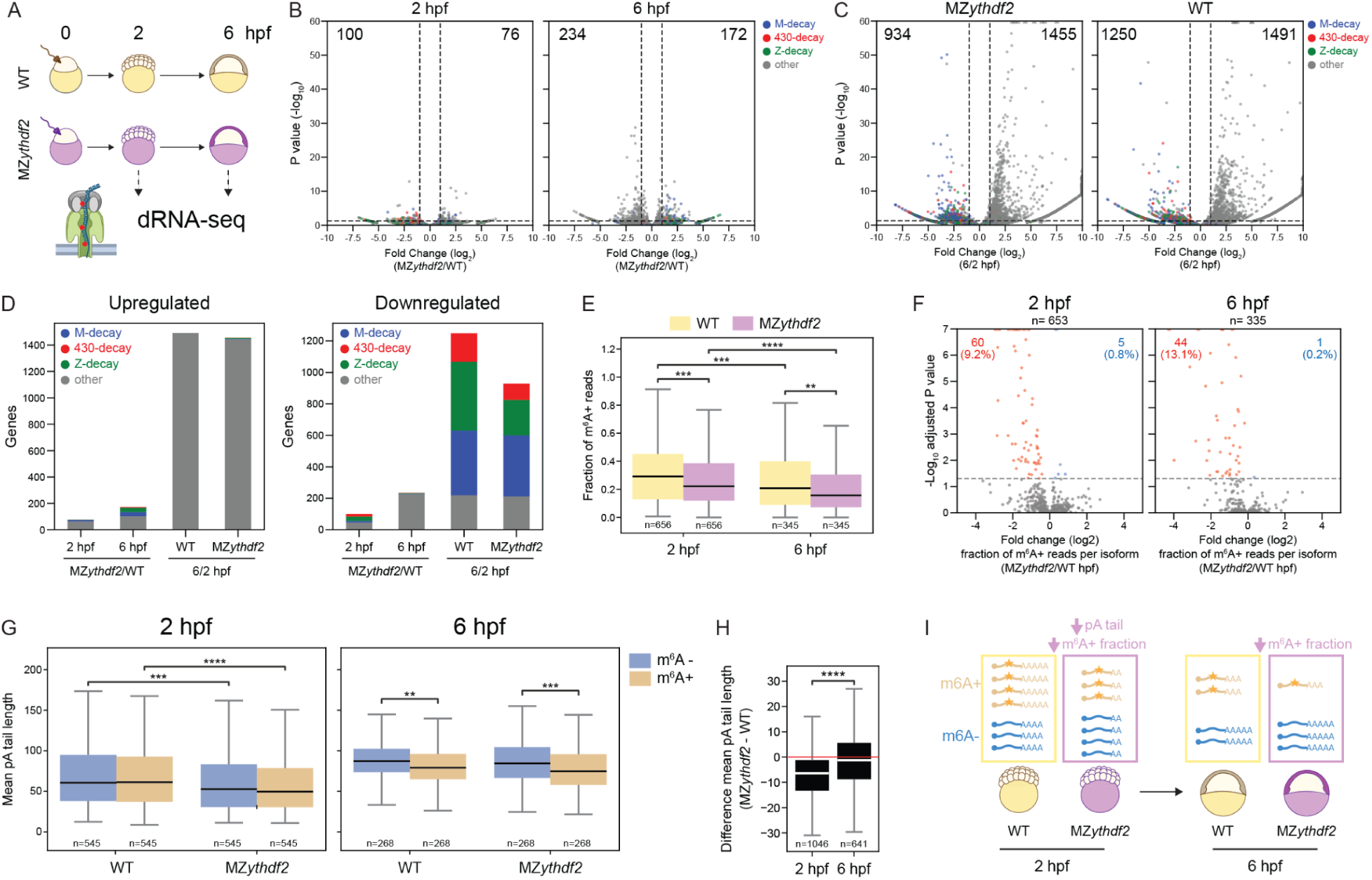
Loss of Ythdf2 does not impair m^6^A-dependent decay and deadenylation but reduces m^6^A stoichiometry and globally shortens poly(A) tails pre-ZGA. (**A**) MZ*ythdf2* mutant and WT sibling embryos were collected at 2 and 6 hpf and sequenced by DRS. (**B**) Volcano plots of differential gene expression between MZ*ythdf2* and WT at 2 hpf (left) and 6 hpf (right), with genes colored by decay mode (M-decay/maternal, blue; zygotic, green; miR-430, red; other, gray). (**C**) Volcano plots of differential gene expression between 2 and 6 hpf for MZ*ythdf2* (left) and WT (right), colored by decay mode as in (B). Points beyond the axis limits are shown as triangles. (**D**) Number of upregulated (left) and downregulated (right) differentially expressed genes in each decay mode, for the MZ*ythdf2*/WT comparison at 2 and 6 hpf and the 6/2 hpf comparison within each genotype. (**E**) Fraction of m^6^A+ reads per methylated isoform in WT (yellow) and MZ*ythdf2* (purple) at 2 and 6 hpf. Groups were compared by Mann-Whitney U test with Benjamini-Hochberg correction (*** P<0.001, **** P<0.0001). (**F**) Volcano plots of the per-isoform difference in m^6^A+ read fraction between MZ*ythdf2* and WT at 2 hpf (left) and 6 hpf (right). Isoforms with a significantly decreased m^6^A+ fraction in MZ*ythdf2* are shown in red (count and percentage, upper left) and those with a significantly increased fraction in blue (upper right). The horizontal dashed line marks adjusted P = 0.05. (**G**) Mean poly(A) tail length of m^6^A- (blue) and m^6^A+ (tan) reads per isoform shared in WT and MZ*ythdf2* at 2 hpf (left) and 6 hpf (right). Statistical significance was assessed by Mann-Whitney U test (** P<0.01, *** P<0.001, **** P<0.0001). (**H**) Difference in mean poly(A) tail length between MZ*ythdf2* and WT for all shared isoforms (methylated, unmethylated, and ambiguous) at 2 hpf and 6 hpf. Statistical significance was assessed by unpaired t-test. (**I**) MZ*ythdf2* embryos exhibit decreased m6A+ read fraction at 2 and 6 hpf and decreased poly(A) tail lengths at 2 hpf compared to WT.

### Loss of Ythdf2 does not impair m^6^A-dependent decay and deadenylation but reduces m^6^A stoichiometry and globally shortens poly(A) tails at the onset of MZT

The extent to which Ythdf2 mediates m^6^A-dependent mRNA fate during MZT remains uncertain^48,49^. To directly examine the contribution of Ythdf2, we performed DRS on maternal-zygotic *ythdf2* (MZ*ythdf2*) and Wild-type (WT) sibling embryos at 2 and 6 hpf (**Fig. 6A**). We first compared expression between MZ*ythdf2* and WT embryos and found that only a small minority of genes were differentially expressed at either time point, most of which did not belong to a defined decay mode (**Fig. 6B, D**). Moreover, while downregulated genes showed no enrichment for methylation status, upregulated genes in MZ*ythdf2* were overrepresented by unmethylated mRNAs (**Supp. Fig. 8A**). The number of genes differentially expressed between 2 and 6 hpf was comparable between MZ*ythdf2* and WT across all decay modes (**Fig. 6C, D**), with downregulated genes enriched for methylated mRNAs and upregulated genes enriched for unmethylated mRNAs in both genotypes (**Supp. Fig. 8B**). These results indicate that loss of Ythdf2 does not affect global maternal mRNA clearance or ZGA, regardless of methylation status. Additionally, the fraction of m^6^A+ reads decreased between 2 and 6 hpf in *MZythdf2* embryos (**Fig. 6E**), indicating that m^6^A-dependent destabilization continues in the absence of Ythdf2. This suggests that the loss of Ythdf2 can be compensated for, consistent with the fact that all three Ythdf paralogs are expressed across MZT (**Supp. Fig. 8C**) and their proposed functional redundancy^49,50^.

Despite the preservation of global and m^6^A-mediated decay, intriguingly, the fraction of m^6^A+ reads per isoform was significantly reduced in MZ*ythdf2* compared to WT at both 2 and 6 hpf (**Fig. 6E, F**). This reduction in m^6^A stoichiometry may suggest the preferential stabilization of m^6^A marked transcripts by Ythdf2 during oogenesis or in the early embryo prior to 2 hpf, such that in its absence fewer m^6^A+ molecules are maternally inherited or maintained.

We next assessed whether loss of Ythdf2 alters the relationship between m^6^A status and poly(A) tail length. As in WT, no significant difference in poly(A) tail length between m^6^A+ and m^6^A- reads was detected at 2 hpf in MZ*ythdf2* embryos (**Fig. 6G**). At 6 hpf, m^6^A+ reads continued to exhibit shorter poly(A) tails than m^6^A- reads in both WT and MZ*ythdf2* embryos (**Fig. 6G**), demonstrating that m^6^A associated poly(A) tail shortening persists independently of Ythdf2 and is likely mediated by redundant reader activity. Strikingly, however, isoforms with m6A in MZ*ythdf2* embryos displayed globally shorter poly(A) tails than their WT counterparts at 2 hpf, regardless of read m^6^A status (**Fig. 6G**, left panel). This m6A-independent shortening was no longer detectable at 6 hpf, where only the difference between m^6^A+ and m^6^A- remained (**Fig. 6G**, right panel). This global poly(A) tail shortening was also apparent when comparing all isoforms (i.e., methylated, unmethylated, and ambiguous), between genotypes at 2 hpf but not 6 hpf (**Fig. 6H; Supp. Fig. 8D**), validating that the effect is m^6^A-independent and specific to the pre-ZGA window. This unexpected finding suggests that Ythdf2 plays a previously unrecognized role in maintaining global poly(A) tail length at the onset of MZT, likely through a mechanism independent of its m^6^A reader function, a role that warrants direct investigation in future work.

Together, these results support a model in which Ythdf2 is not the primary mediator of m^6^A-dependent maternal mRNA decay and deadenylation during MZT, consistent with the genetic evidence from Kontur *et al.*^49^. Nevertheless, our data reveal previously unrecognized contributions of Ythdf2 to m^6^A stoichiometry maintenance and global poly(A) tail homeostasis at the earliest stages of embryonic development (**Fig. 6I**). Unlike its role in maternal mRNA decay, these functions are not compensated by Ythdf1 or Ythdf3. The single-molecule resolution of DRS makes it possible to disentangle these distinct roles of Ythdf2 from the confounding effects of transcript sequence and isoform identity that obscure such relationships in short-read sequencing data.

## Discussion

By applying nanopore DRS to a developmental time course in zebrafish embryos, we generated the first single-molecule, multi-feature epitranscriptomic atlas of vertebrate MZT. We simultaneously resolved m^6^A modification status, poly(A) tail length, and isoform identity on each sequenced RNA molecule, capabilities inaccessible to antibody-based bulk methods. Whereas prior DRS-based m^6^A studies have been conducted predominantly in steady-state systems^59,60,72–75^, the dynamic transcriptomic remodeling of MZT provides a uniquely informative system for dissecting the functional consequences of m^6^A on transcript fate at single-molecule resolution (**Fig. 1A**). This resolution revealed that the zebrafish maternal transcriptome is far more broadly methylated than previously estimated, with more than 78% of expressed genes harboring methylated isoforms at 2 hpf compared to prior bulk estimates of one-third^48^, likely reflecting the improved sensitivity of DRS at low-stoichiometry sites below the detection threshold of IP-based methods.

Unmethylated genes display a structural predisposition against methylation: shorter transcript length, fewer EJC-distal DRACH motifs, and shorter exons (**Fig. 1D**, **Supp. Fig. 1D,E**). These genes also exhibit functional enrichment for core housekeeping machinery (**Fig. 1G**). By contrast, methylated genes encode developmental regulators and signaling proteins (**Fig. 1F**). Together, these observations suggest that m^6^A deposition selectively marks transcripts requiring spatiotemporal control during MZT. A similar functional dichotomy has been reported in human preimplantation embryos^76^, suggesting this selectivity is a conserved feature of vertebrate MZT.

A central advance of this study is the demonstration that m^6^A promotes maternal transcript decay and poly(A) tail shortening independent of sequence, established by directly comparing m^6^A+ and m^6^A- reads within the same isoform. This within-isoform analysis controls for all sequence-intrinsic features, such as codon usage and 5’- and 3’-UTR composition, that confound cross-transcript comparisons in short-read datasets. The m^6^A+ read fraction of individual isoforms declined progressively across MZT (**Fig. 3C,E**, **4B**, **Supp. Fig. 4A**), and this decline was preserved in strictly maternal genes without detectable zygotic contribution (**Fig. 3D**, **Supp. Fig. 4A**, **Supp. Fig. 5A**). This indicates that the decreasing proportion of methylated molecules reflects preferential decay of m^6^A marked transcripts rather than dilution by newly synthesized unmethylated zygotic mRNAs. At the poly(A) tail level, m^6^A+ reads show comparable tail lengths to their m^6^A- counterparts at 2 and 4 hpf but significantly shorter tails at 6 hpf (**Fig. 3G,H**), directly associating m^6^A with poly(A) tail shortening at the single-molecule level (**Fig. 3I, Supp. Fig. 4C-E**). These results provide direct single-molecule evidence for the m^6^A-deadenylation relationship previously inferred from population-level data in zebrafish^49^.

Our positional analysis reveals a striking temporal divergence in how m^6^A location shapes its effect on poly(A) tail length, a previously undescribed dimension of m^6^A regulation during vertebrate development (**Fig. 4H**). CDS m^6^A is constitutively associated with decay and shorter poly(A) tails from 2 hpf, before ZGA (**Fig. 4B-G**). This is consistent with the CDS-m^6^A decay (CMD) pathway in which ribosome pausing at m^6^A-modified codons triggers rapid, translation-coupled transcript destabilization^39–41,77^. The progressive decline in CDS m^6^A+ reads across MZT, coinciding with rising global translation rates during this period^2,3,7^, supports a model in which increasing ribosome engagement accelerates CMD-mediated clearance of CDS-methylated transcripts. By contrast, 3′-UTR m^6^A shows a mixed, bidirectional effect on poly(A) tail length at 2 hpf (**Fig. 4C**) and acquires net poly(A)-shortening activity only at 6 hpf (**Fig. 4C-G**). This temporal shift aligns with the observation that all three Ythdf paralogs associate with maternal mRNAs in a ZGA-dependent manner^10^. This provides a mechanistic basis for the absence of 3’-UTR m^6^A mediated deadenylation before ZGA, which emerges only as Ythdf proteins become competent to engage their targets and recruit CCR4-NOT deadenylase activity. The subset of isoforms showing longer tails in m^6^A+ reads at 2 hpf may instead reflect the pre-ZGA stabilizing activity of Igf2bp3, which maintains m^6^A-bound maternal transcripts in a translationally competent state before Ythdf engagement^42^.

Together, these findings reveal two novel principles of m^6^A regulation during MZT. First, the same mRNA sequence can follow different temporal fate trajectories depending solely on where m^6^A is deposited, with CDS m^6^A driving poly(A) tail shortening and repression throughout MZT while 3′-UTR m^6^A permits early expression before becoming destabilizing after ZGA (**Fig. 4C-H**). Second, this temporal divergence implies that CDS and 3′-UTR m^6^A engage distinct effector machinery. CDS m^6^A acts before ZGA, when Ythdf proteins are not yet bound to mRNA in zebrafish^10^, indicating that its effect operates through a translation-coupled mechanism likely independent of Ythdf readers. In contrast, 3’-UTR m^6^A activity emerges only after ZGA, consistent with the timing of Ythdf engagement and recruitment of the CCR4-NOT deadenylase complex.

The finding that m^6^A positional context determines transcript fate raises the question of what governs where m^6^A is deposited in the first place. Our machine learning analysis confirms that the same architectural rules established in somatic systems operate in the vertebrate embryo (**Fig. 5**)^23,24,78,79^. EJC-mediated occlusion of METTL3 access near splice junctions suppresses methylation across much of the CDS, while stop codon-proximal DRACH motifs in long terminal exons are the most efficiently modified sites, a topology that directly explains why 3′-UTR m^6^A predominates in the maternal transcriptome (**Fig. 1B and 4A**)^34–36^. That these determinants remain stable across the entire MZT time course suggests that the deposition machinery operates through fixed structural rules rather than being dynamically remodeled in response to developmental signals. Beyond architecture, flanking nucleotides at positions −4 and +4 relative to DRACH contribute independently to stoichiometry prediction (**Fig. 5F,G** and **Supp. Fig. 7B-D**), suggesting METTL3 recognition extends beyond the minimal five-nucleotide motif. GGACU and AGACU are the most frequently modified DRACH variants (**Fig. 5B** and **Supp. Fig. 6A-C and 7A**), consistent with prior zebrafish profiling^48^ and structural features favoring METTL3 engagement.

The contested role of Ythdf2 in zebrafish MZT has been difficult to resolve with population-level sequencing data, given the inability of short-read bulk methods to link modification state directly to transcript fate^48,49,80^. Our single-molecule analysis of MZ*ythdf2* embryos both confirms and extends the current genetic understanding, revealing previously unrecognized contributions of this reader to m^6^A stoichiometry and poly(A) tail homeostasis. Consistent with the genetic evidence from Kontur *et al.*^49^, global maternal mRNA clearance, ZGA, and decay mode dynamics are largely unaffected by loss of Ythdf2 (**Fig. 6B-D**). This conclusion is further reinforced by the persistence of m^6^A-associated decay and poly(A) tail shortening in MZ*ythdf2* embryos at 6 hpf compared to 2 hpf (**Fig. 6E,G** and **Supp. Fig. 8B**), which we attribute to redundant activity of Ythdf1 and Ythdf3, both of which engage maternal mRNAs post-ZGA^50^.

Two findings, however, reveal novel roles for Ythdf2 in this developmental context that are not compensated by the other Ythdf paralogs. First, the fraction of m^6^A+ reads per isoform is significantly reduced in MZ*ythdf2* embryos at both 2 and 6 hpf (**Fig. 6E,F**). This is consistent with a model in which Ythdf2 preferentially stabilizes m^6^A marked transcripts during oogenesis or in the early embryo prior to 2 hpf. In its absence, fewer m^6^A+ molecules are maternally deposited or maintained at the onset of MZT. A role for Ythdf2 in shaping the maternal transcriptome is also supported by evidence from mice, where YTHDF2 loss in oocytes causes female infertility and dysregulates the maternal transcriptome^46^. More broadly, this is consistent with the principle that m^6^A reader function is context-dependent, with the same reader producing distinct effects in different cellular environments^81^. Second, transcripts in MZ*ythdf2* embryos display globally shorter poly(A) tails at 2 hpf regardless of m^6^A status, an effect that resolves completely by 6 hpf (**Fig. 6G,H; Supp. Fig. 8D**). This pre-ZGA, m^6^A-independent reduction in global poly(A) tail length is unexpected and suggests that Ythdf2 plays a role in maintaining poly(A) tail homeostasis at the earliest stages of embryonic development likely through a mechanism distinct from its m^6^A-reader function. In this regard, the milder developmental consequence of Ythdf2 loss in zebrafish compared to mouse may reflect the broader cytoplasmic polyadenylation activity characteristic of zebrafish embryos, which is driven by the near-equal expression of CPEB1 and CPEB4 and recognition of a more permissive CPE motif than in other vertebrates^14^. We speculate that this enhanced re-adenylation capacity may partially compensate for the globally shorter poly(A) tails observed in MZ*ythdf2* zebrafish embryos, buffering the developmental consequences of Ythdf2 loss in a way that the more restricted cytoplasmic polyadenylation activity in mouse cannot.

A limitation of our approach is that DRS requires the presence of a terminal poly(A) tail for adapter hybridization and library capture, meaning that fully deadenylated or terminally uridylated transcripts are not recovered. While the majority of deadenylated and uridylated maternal mRNAs are subsequently degraded during MZT^8,13,21,82^, we cannot exclude that a small fraction may be stabilized in a deadenylated or uridylated form, in which case their absence from our dataset would be interpreted as decay rather than stable accumulation in a tail-shortened state. A second caveat concerns our MZ*ythdf2* analysis. Whether the observed changes reflect direct Ythdf2 binding to affected transcripts or secondary consequences of Ythdf2-mediated regulation of key upstream regulators cannot be resolved from our data alone. Addressing this will require integration of transcriptome-wide Ythdf2 binding data with single-molecule DRS profiling of MZ*ythdf2* and WT oocytes and embryos at earlier developmental stages.

## Methods

### Zebrafish maintenance

Wild-type zebrafish (*Danio rerio*) embryos were obtained through natural mating of the TU-AB strain of mixed ages (5–18 months). Mating pairs were randomly chosen from a pool of 60 males and 60 females allocated for each day of the month. MZ*ythdf2* has an 8 bp deletion in the *ythdf2* genes as described previously^48,49^, and were compared to Wild-type siblings. Embryos and adult fish were maintained at 28 °C. Fish lines were maintained according to the International Association for Assessment and Accreditation of Laboratory Animal Care research guidelines, and protocols were approved by the Yale University Institutional Animal Care and Use Committee (IACUC).

### Embryos injection

All injections into zebrafish embryos were performed on dechorionated one-cell stage embryos with 1 nL volumes. To inhibit RNA Polymerase II, embryos were injected with 0.2 ng of α-amanitin (Sigma-Aldrich, A2263) resuspended in nuclease-free water. To inhibit miR-430 activity, embryos were injected with 10 µM of tiny locked nucleic acid (tinyLNA), complementary to the seed region of miR-430 (5′-TAGCACTT-3′ (Exiqon)).

### Sample collection and purification

For direct RNA sequencing with nanopore, 25 embryos at the specified stage were transferred to 1.5-mL tubes and flash frozen in liquid nitrogen. Frozen embryos were thawed and actively lysed with 1 mL of TRIzol (Invitrogen), and total RNA was extracted according to the manufacturer’s protocol. Poly(A)+ mRNAs were isolated with oligo d(T)_25_ magnetic beads (New England BioLabs) according to the manufacturer’s protocol and eluted in 30 µL of water.

### DRS library preparation and sequencing

Poly(A)+-selected RNA samples were prepared for nanopore sequencing using the direct RNA sequencing kit (SQK-RNA002), following ONT protocol version DRS_9080_v2_revI_14Aug2019 with half reactions for each library up to the RNA Adapter (RMX) ligation step, with the following adaptations. Briefly, each half reaction started with 250 ng of poly(A)+ RNA. Samples were ligated to pre-annealed custom RT adaptors (IDT) containing barcodes^83^ using concentrated T4 DNA Ligase (NEB-M0202M) for 15 min at room temperature. For non-barcoded runs, the commercial RT adapter included in the SQK-RNA002 kit was used instead. Reverse transcription was then performed for 15 min at 50 °C using SuperScript IV reverse transcriptase (Invitrogen,18090050), followed by heat inactivation for 5 min at 70 °C. Ligated products were purified using 1.8X Agencourt RNAClean XP beads (Fisher Scientific, NC0068576) and washed with freshly prepared 70% ethanol. For the final ligation, 50 ng of reverse-transcribed RNA from each reaction was pooled, mixed with RMX adapter (sequencing adapter bearing the motor protein), and incubated for 15 min with concentrated T4 DNA Ligase (NEB-M0202M). For non-barcoded runs, 200 ng of reverse-transcribed RNA was used for ligation with the RMX adapter. The ligated RNA:DNA hybrid was purified using 1X Agencourt RNAClean XP beads and washed twice with Wash Buffer (WSB). Samples were eluted in Elution Buffer (EB), mixed with RNA Running Buffer (RRB), and loaded onto a primed R9.4.1 flowcell on a MinION or PromethION device.

### Basecalling and demultiplexing direct RNA sequencing data

Raw Fast5 files from DRS sequencing runs were processed with *Master of Pores* pipeline^84^ version 2 (https://github.com/biocorecrg/MOP2). The *mop_preprocess* module was used to basecall the raw signal with *Guppy* v6.0.6 (Oxford Nanopore Technologies) using the m^6^A-aware basecalling model *rna_r9.4.1_70bps_m6A_hac.cfg* (m^6^ABasecaller)^59^, available at https://github.com/novoalab/m6ABasecaller/tree/main/basecalling_model. If the run was multiplexed (see **Table S1**), the *DeePlexiCon* algorithm^83^ was used to demultiplex the reads.

### Extraction of modification information from DRS datasets basecalled with the m^6^A basecaller

m^6^A-basecalled Fast5 files were processed with *ModPhred*^85^ (https://modphred.readthedocs.io/en/latest/) to encode the modification probabilities computed by the *m^6^ABasecaller* into FASTQ files. *ModPhred* was then used to map these reads to the GRCz11 genome with minimap2 2.17-r941^86^ using the parameters “*-ax map-ont -k13”*, and to generate a compressed bedMethyl (*.mod.gz) file listing all m^6^A sites, coverage, and modification frequency (stoichiometry). The modification frequency is calculated as the percentage of reads with m6A at that genome position.

### Gene and isoform annotation for individual reads

Per-read gene and isoform annotation was performed on all unique and primary alignments in each replicate, extracted using the samtools flag “*-F 3488*”, with *Isoquant* v3.3.0^87^ using parameters *“--complete_genedb --data_type nanopore*” and the zebrafish Ensembl annotation version 110. Isoquant assigns reads to isoforms based on intron-chain matching and exonic overlap detection.

### Poly(A) tail length estimation

Per-read poly(A) tail length estimation was performed using the *mop_tail* module of MoP2^84^ with *tailfindr* v1.3^88^. Reads returning NaN values were excluded from downstream analyses.

### Identification of replicable m^6^A-modified sites

The *ModPhred* output file (*.*mod.gz*) was processed with a custom Python script (https://github.com/novoalab/m6ABasecaller_per_read_analysis/blob/main/PerPosition_Analysis/ModPhred_PostProcessing.py) to identify replicable m^6^A sites. Replicable sites were defined as those with coverage ≥25 and a modification frequency ≥5% in all replicates of a given condition. Requiring replication within, rather than across, conditions accounts for the decay of maternal transcripts over the time course while limiting false positives. Only m^6^A calls at these whitelisted replicable sites were considered for all downstream per-read analyses. Replicable site information was compiled into Supplemental Dataset 8.

### Generation of a per-read table with m6A, poly(A) tail and gene information

A nextflow-based pipeline (*m6AB_PerRead*; https://github.com/novoalab/m6ABasecaller_per_read_analysis/) was developed to generate tables containing m^6^A-modified sites, poly(A) tail length estimates, and gene and isoform assignments for each individual read. The pipeline takes three inputs: the *ModPhred* alignment files, pre-filtered to unique and primary alignments using the samtools flag “*-F 3488*”; the poly(A) tail length estimates from the *mop_tail* module of MoP2; and the read-to-gene/isoform assignment from IsoQuant. From the alignment file, a custom script extracts the modified positions in each read, if any, along with additional features including the number of modified positions, alignment start and end coordinates, strand and total read length. This table is then filtered to retain only replicable modified sites (defined above) and merged with the poly(A) tail and gene assignment data.

### Gene-level analysis of MZT time course samples

Using the per-read table, reads that did not map uniquely to a single isoform (transcript ID) were removed, as were isoforms with fewer than 25 reads in each replicate and reads lacking a *tailfindr* poly(A) tail estimate (NaN values). Each read was then assigned to a gene ID based on its transcript ID. Genes were then classified by their number of m^6^A+ reads: those with no m^6^A+ reads were labeled *Unmethylated*, those with 1-3 m^6^A+ reads *Ambiguous*, and those with ≥4 *Methylated*.

### Gene Ontology (GO) analysis

GO analysis was performed and plotted using custom Python scripts. Zebrafish GO term sets: (GO_Molecular_Function_2018, GO_Cellular_Component_2018, GO_Biological_Process_2018, and WikiPathways_2018) were retrieved from the Enrichr database using the *GSEApy* module. For each analysis, a list of genes of interest (e.g., upregulated or downregulated) was compiled alongside a background list of all genes that were expressed and passed filtering in the relevant condition. Fold enrichment was calculated as the fraction of genes of interest annotated to a given ontology term divided by the corresponding fraction of background genes.

### Differential expression

Using the per-read table, reads that did not map uniquely to a single gene (gene ID) were removed and counts were summed to obtain a per-gene read count per replicate (**Supp. Dataset 3, 4**). Differential expression analysis was then performed on these per-gene read counts using DESeq2 v1.40.1^89^. For the WT time course analysis, genes were required to have at least 100 reads summed across samples, whereas for the MZ*ythdf2* and corresponding WT sibling analysis, a lower threshold of 10 reads was applied owing to the reduced global read depth. Genes with an absolute log_2_ fold change greater or equal to 1.0 and an adjusted P-value lower or equal than 0.05 were considered differentially expressed.

### Isoform-level and read-level analysis of MZT time course samples

The following filtering was used for all isoform-level analyses unless otherwise stated. Using the per-read table, reads that did not map uniquely to a single isoform (transcript ID) were removed, as were isoforms with fewer than 25 reads in each replicate and reads lacking a *tailfindr* poly(A) tail estimate (NaN values) (**Supp. Dataset 5**). Reads from each replicate were then combined. Isoforms were classified by their number of m^6^A+ reads as *Unmethylated* (0), *Ambiguous* (1-3), or *Methylated* (≥4). To characterize expression from an external dataset, ribosomal RNA-depleted RPKM values were obtained from Vejnar *et al.* 2019^9^, after filtering for isoforms with >5 RPKM. Additionally, poly(A)-selected RPKM values were obtained from this dataset for Ythdf paralog expression over time. Translation efficiency values were obtained from Beaudoin *et al.* 2018^67^. DRS poly(A) tail lengths were compared to PAL-seq data tail lengths with ≥50 PAL-seq tags and non-NaN RNA-seq values from Subtelny *et al.*^15^. Unless otherwise stated in the figure legends, Mann-Whitney tests were used to assess differences between groups. P values for differences in regression slopes (b1 and b2) were calculated as 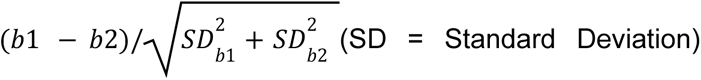, and the resulting t statistic was evaluated against the Student’s t distribution^90^. For poly(A) tail length analysis of α-amanitin and miR-430 LNA-treated samples, the same filtering strategy was applied, except that the minimum number of reads per replicate was lowered from 25 to 10 to accommodate the lower sequencing depth (**Supp. Dataset 6**).

### Categorization of decay mode pathways

Gene-level decay mode classifications were obtained from Vejnar *et al.* 2019^9^, and isoforms were assigned the decay mode of their corresponding gene. Genes or isoforms not classified in the Vejnar *et al.* dataset were labeled as “other”.

### DRACH motif methylation analysis of MZT time course samples

Only uniquely mapping reads were used and merged with zebrafish Ensembl 110 transcript annotations. Read start and end genomic coordinates and m^6^A-modified sites were mapped onto transcript annotation using custom Python scripts, and the portion of the annotated sequence covered by each read was extracted. Within each extracted sequence, DRACH motifs were identified using the regular expression (DRACH=([AGT][AG]AC[ACT])), and each motif was assessed for the presence of m^6^A at the adenosine position. DRACH motifs, their flanking nucleotides, and their m^6^A modification status were compiled, and position weight matrices of methylated sites were used to generate motif logos at each timepoint.

### Machine learning analysis of per site stoichiometry

For machine learning analyses, only replicable m^6^A modified sites that mapped to single isoform (transcript ID) at all timepoints (2, 4, and 6 hpf) and their corresponding site modification frequencies were used. For feature-importance analysis of per-site stoichiometry, features were derived from zebrafish Ensembl 110 annotations. The features investigated were: m^6^A site position within the transcript; distance to closest splice site; distance to left and right splice site; number of exons in the transcript; presence within a DRACH motif (True/False); exon position from the transcript start and from the end; kmer (1- to 5-mer) counts of the left and right flanking sequence (75 nt each); the transcript region in which the site falls (5’-UTR, CDS, or 3’-UTR); 5’-UTR and 3’-UTR AT and GC content; transcript length; minimum replicate coverage; distance from the start and stop codons; and the identity of the DRACH motif containing the modification. For each stoichiometry module (2, 4, and 6 hpf), features were split into training and validation sets at a 80:20 ratio, with the training set used for hyperparameter tuning. An eXtreme Gradient Boosting (XGBoost) model was trained with hyperparameter tuning by halving grid search and 10-fold cross-validation. Model interpretation was performed using SHapley Additive exPlanations (SHAP) via the *shap* library, with the *TreeExplainer* module generating a SHAP matrix matching the dimensions of the input features. Global feature importance was calculated as the mean of the absolute values of this matrix.

For sequence-importance analysis of per-site stoichiometry, transcript sequences spanning -20 to +20 nucleotides around each filtered m^6^A site were extracted from the Ensembl annotation and one-hot encoded into a position × nucleotide matrix. An XGBoost model was then trained with hyperparameter tuning by halving grid search and 10-fold cross-validation, with nucleotide positions split into training and validating sets at an 80:20 ratio. Global feature importance was again calculated as the mean of the absolute values of the SHAP matrices generated with the *TreeExplainer* module.

### MZ*ythdf2* and WT sibling comparison analysis

Using the per-read table, reads that did not map uniquely to a single isoform (transcript ID) were removed, as were isoforms with fewer than 25 reads in each replicate and reads lacking a *tailfindr* poly(A) tail estimate (NaN values) (**Supp. Dataset 7**). Reads from each replicate were then combined. Decay modes were assigned as described above. The fraction of m^6^A+ reads was calculated for methylated isoforms shared between *MZythdf2* and WT sibling at each timepoint. Per-isoform poly(A) tail length was calculated as the mean of *tailfindr* values across reads (m^6^A- or m^6^A+) of that isoform. Mann-Whitney tests were used for statistical comparisons unless otherwise stated in figure legends.

## Data availability

Raw sequencing data generated in this study have been deposited in the European Nucleotide Archive (ENA) database, under accession codes compiled in **Table S1**. Output tables from m^6^ABasecaller pipeline, filtered data tables, and processed count tables are included as Supplementary Datasets. Additional data supporting the findings of this study are available from the corresponding author upon request.

## Code availability

Code to process nanopore sequencing data basecalled with m^6^ABasecaller has been made publicly available in GitHub (https://github.com/novoalab/m6ABasecaller_per_read_analysis). All scripts used to process and analyze DRS data will be released at the time of publication in Github.

## Declaration of generative AI and AI-assisted technology use

During the preparation of this work, the authors used Claude and ChatGPT to improve text readability and style and to optimize code. After using these tools, the authors reviewed and edited all text and code. The authors take full responsibility for the content of this publication.

## Acknowledgements

We thank all members of the Beaudoin lab for feedback and support. AD-T was supported by an FPI Severo-Ochoa fellowship by the Spanish Ministry of Economy, Industry and Competitiveness (MEIC). This work was supported by the Australian Research Council (DP180103571 to EMN), by the Spanish Ministry of Economy, Industry and Competitiveness (MEIC) (PID2021-128193NB-100 to EMN) and the European Research Council (ERC-StG-2021 No 101042103 to EMN). We acknowledge the support of the MEIC to the EMBL partnership, Centro de Excelencia Severo Ochoa and CERCA Programme / Generalitat de Catalunya. SAA was supported by an Ruth L. Kirschstein National Research Service Award (NRSA) Individual Predoctoral Fellowship (F31HD114435). JDB was supported by the National Institute of Health (NIH) Maximizing Investigators’ Research Award (MIRA) (R35GM146883).

## Contributions

JDB and EMN conceived the project, acquired funding, and supervised the project. CK and JDB collected RNA samples and performed poly(A)-selection. LLL and RM performed nanopore direct RNA sequencing library preparation, together with ADT (MiNION flowcell runs). ADT performed bioinformatic nanopore raw data processing, including basecalling with m6Basecaller model, transcript assignment, predictions of poly(A) tail lengths, and generation of per-read and per-position data tables. SAA and SM analyzed data and produced data visualizations. SM performed machine learning analysis. SAA, SM, and JDB interpreted the data. SAA and JDB wrote the manuscript with input and approval from all authors.

## Summary of supplementary table and datasets (Supp. Datasets available upon request prior to publication)

| Name | Description |
| --- | --- |
| Supp. Table S1 | Names of samples sequenced, their sequencing details and database information, and raw read count. |
| Supp. Dataset 1 | For each isoform, for each timepoint m<sup>6</sup>A+ and m<sup>6</sup>A- # of reads and m<sup>6</sup>A+ fraction of reads. |
| Supp. Dataset 2 | Tables of gene level m<sup>6</sup>A+ fraction and their cumulative fraction rank for each timepoint |
| Supp. Dataset 3 | Count tables used for DESeq2 analysis Wildtype Untreated Timecourse all timepoints |
| Supp. Dataset 4 | Count tables used for DESeq2 analysis *MZythdf2* and sibling WT all timepoints |
| Supp. Dataset 5 | Filtered PerRead table for WT timecourse 2hph r1,r2; 4hpf r1, r2; 6hpf r1,r2 |
| Supp. Dataset 6 | Filtered PerRead table for α-amanitin r1,r2; LNA r1,r2 |
| Supp. Dataset 7 | Filtered PerRead table for *MZythdf2* r1,r2;siblingWT r1,r2 at 2hpf and 6hpf |
| Supp. Dataset 8 | Raw Per-Position Replicable site table |
| Supp. Dataset 9 | Per-Read tables from m<sup>6</sup>ABasecaller pipeline output from untreated WT samples |
| Supp. Dataset 10 | Per-Read tables from m<sup>6</sup>ABasecaller pipeline output from α-amanitin and LNA samples |
| Supp. Dataset 11 | Per-Read tables from m<sup>6</sup>ABasecaller pipeline output from MZy<sup>thdf2</sup> and sibling WT samples |

---

**Supplementary Figure 1.**
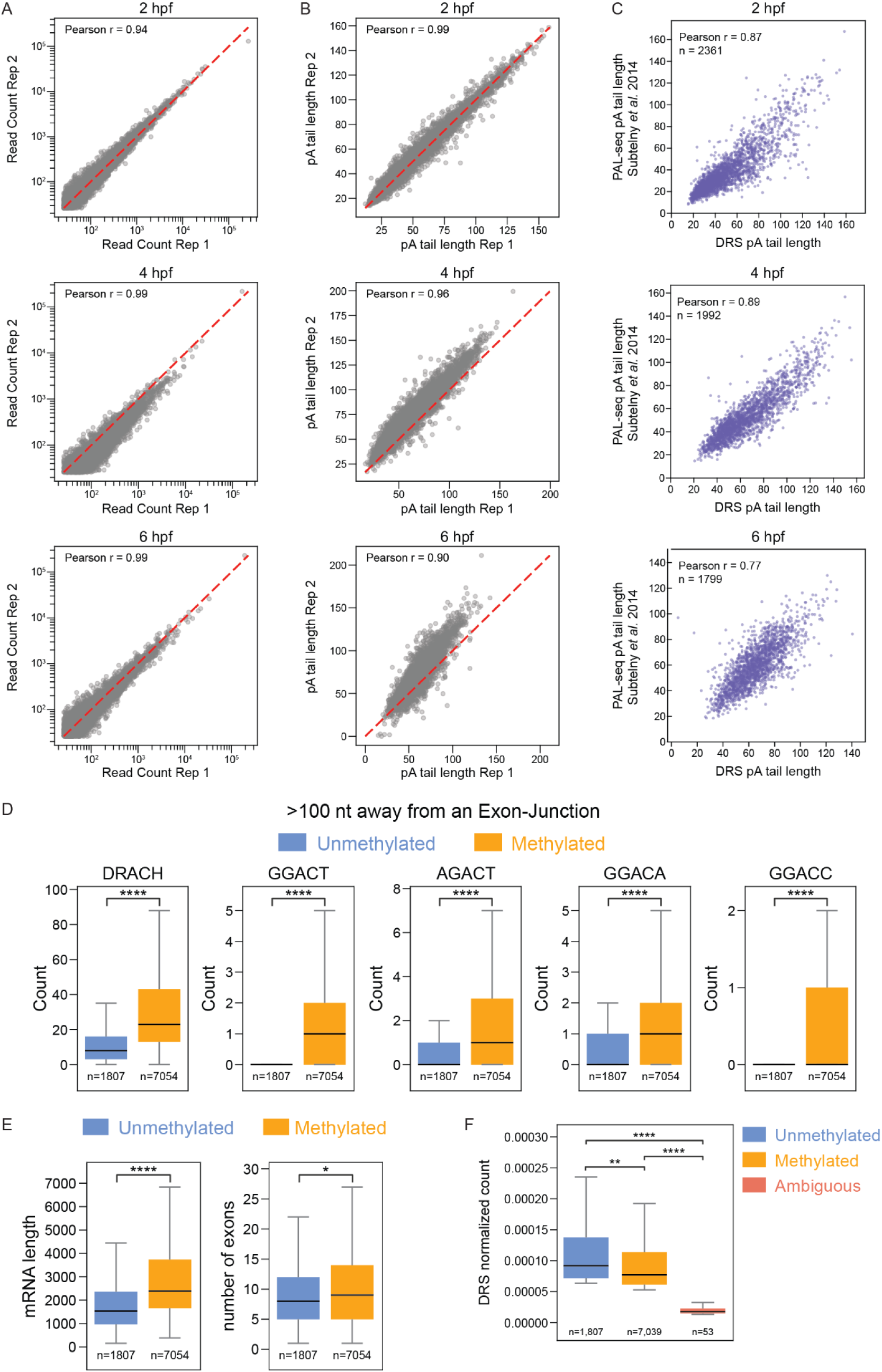
Reproducibility of DRS replicates and poly(A) tail measurements and the structural predisposition of unmethylated isoforms against methylation. (**A,B**) Scatter plots comparing per-isoform read counts (A) and mean poly(A) tail length (B) between biological replicates at 2 hpf (upper), 4 hpf (middle), and 6 hpf (lower). The red dashed line indicates y = x; Pearson *r* is shown for each comparison. (**C**) Scatter plot comparing mean poly(A) tail lengths per isoform from Subtelny *et al.* PAL-seq data^15^ and our DRS data at 2 hpf (upper), 4 hpf (middle), and 6 hpf (lower). (**D**) Counts of all DRACH motifs and of specific DRACH variants in the mRNA sequence, excluding the 100-nucleotide regions flanking exon-exon junctions, for unmethylated and methylated isoforms. (**E**) Annotated mRNA length and number of exons for unmethylated and methylated isoforms. (**F**)DRS read counts per isoform, normalized to total reads per condition, for unmethylated, methylated, and ambiguous isoforms at 2 hpf. n values are indicated below each box. Statistical significance was assessed by Mann-Whitney test (* P<0.05, ** P<0.01, *** P<0.001, **** P<0.0001).

**Supplementary Figure 2.**
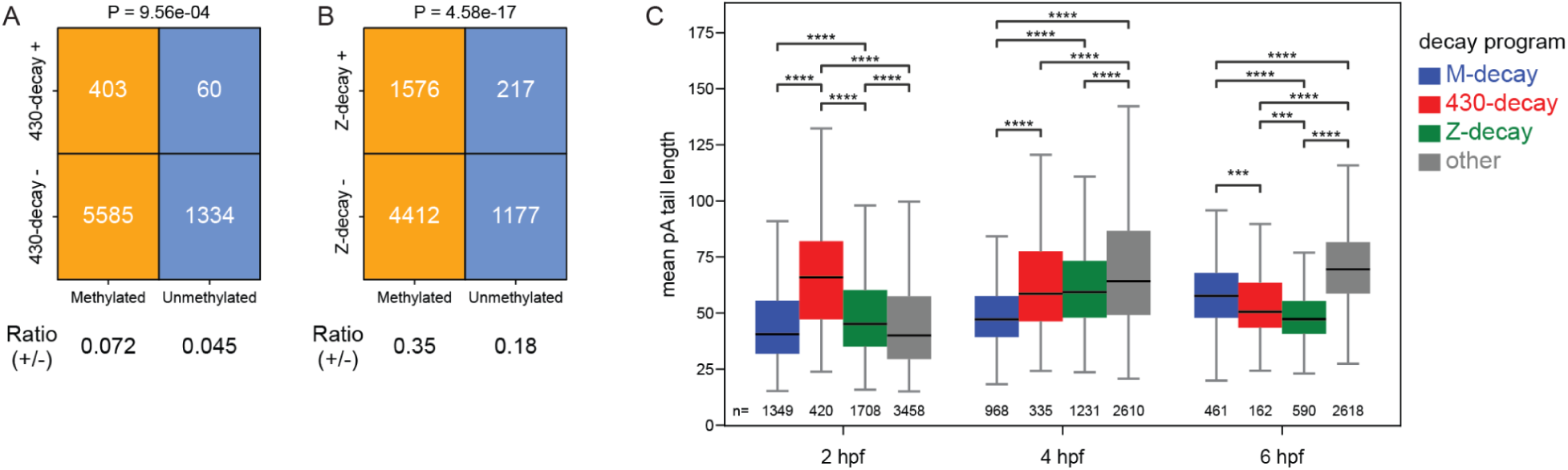
m^6^A enrichment across decay pathways and decay-mode-specific poly(A) tail length across MZT. (**A,B**) Contingency tables of methylated and unmethylated isoforms within versus outside the miR-430 (430-decay) decay (A) and zygotic (Z-decay) (B) pathways at 2 hpf. The *P*-values were calculated by the chi-square test of independence. (**C**) Mean poly(A) tail length of isoforms in each decay program (maternal/M-decay, blue; miR-430/430-decay, red; zygotic/Z-decay, green; other, gray) at 2, 4, and 6 hpf. *n* values are indicated below each box. Statistical significance was assessed by Mann-Whitney test with Benjamini-Hochberg correction (* P<0.05, ** P<0.01, *** P<0.001, **** P<0.0001).

**Supplementary Figure 3.**
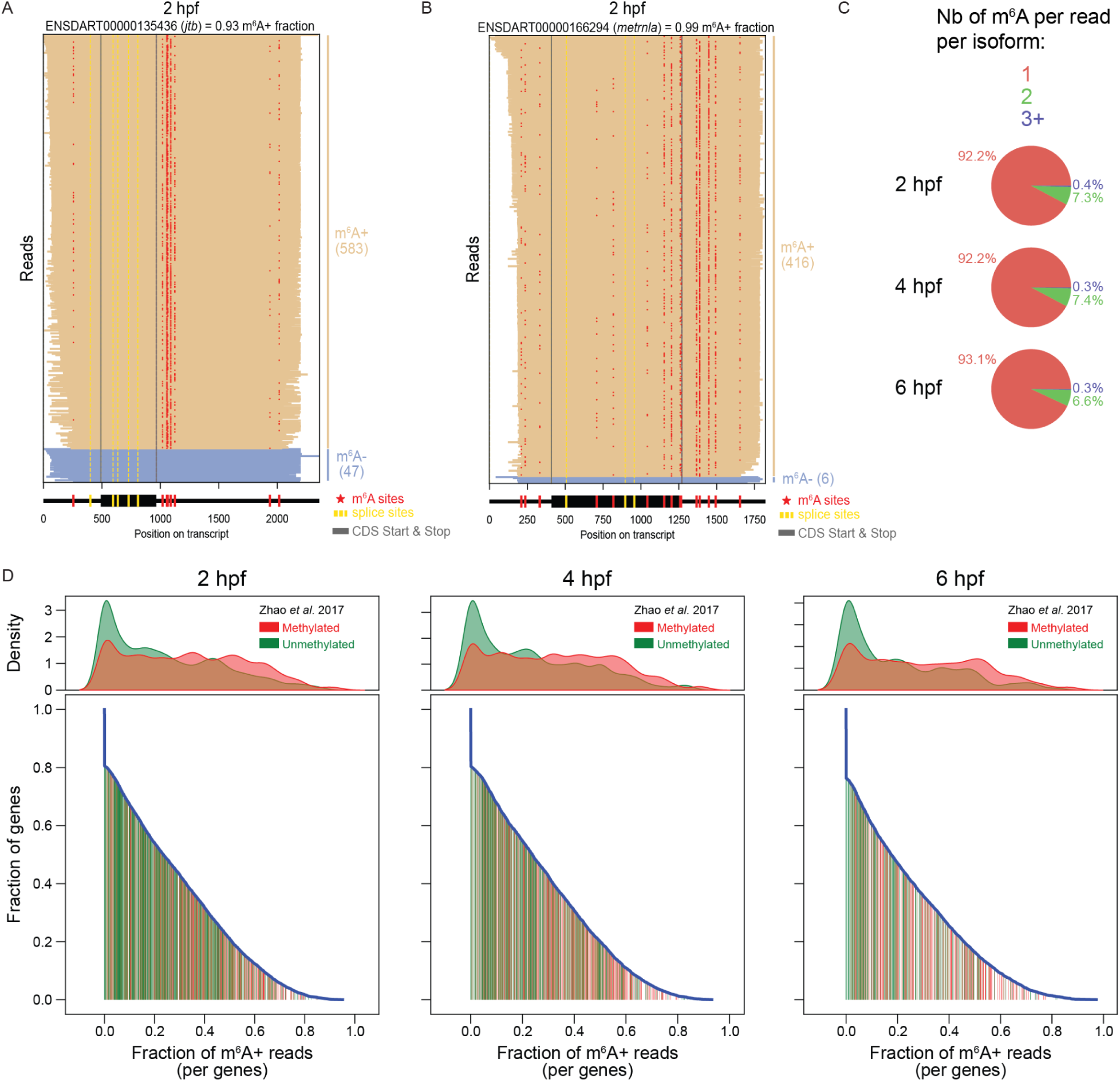
Read-level m^6^A profiles, per-read number of m^6^A modifications, and concordance of DRS m^6^A+ fractions with published antibody-based classifications. (**A**) Read-level view of a representative isoform (ENSDART00000135436, *ilf3*) with a high m^6^A+ fraction (0.93) at 2 hpf. Each row represents a single read; m^6^A sites (red), splice sites (yellow), and CDS start and stop (gray) are indicated below, with m^6^A+ and m^6^A- read counts shown at right. (**B**) Read-level view of a second representative isoform (ENSDART00000166294, *metrnl*) with a high m^6^A+ fraction (0.99) at 2 hpf, displayed as in (A). (**C**) Distribution of the number of m^6^A modifications per read per isoform containing m^6^A at 2, 4, and 6 hpf. (**D**) Upper: kernel density estimate of the per-gene m^6^A+ read fraction in the DRS dataset for isoforms classified as methylated (red) or unmethylated (green) by Zhao *et al.*^48^ Lower: cumulative fraction of genes with an m^6^A+ read fraction greater than or equal to a given value (blue curve), with individual Zhao *et al.* methylated (red) and unmethylated (green) genes shown as vertical lines beneath the curve, at 2, 4, and 6 hpf.

**Supplementary Figure 4.**
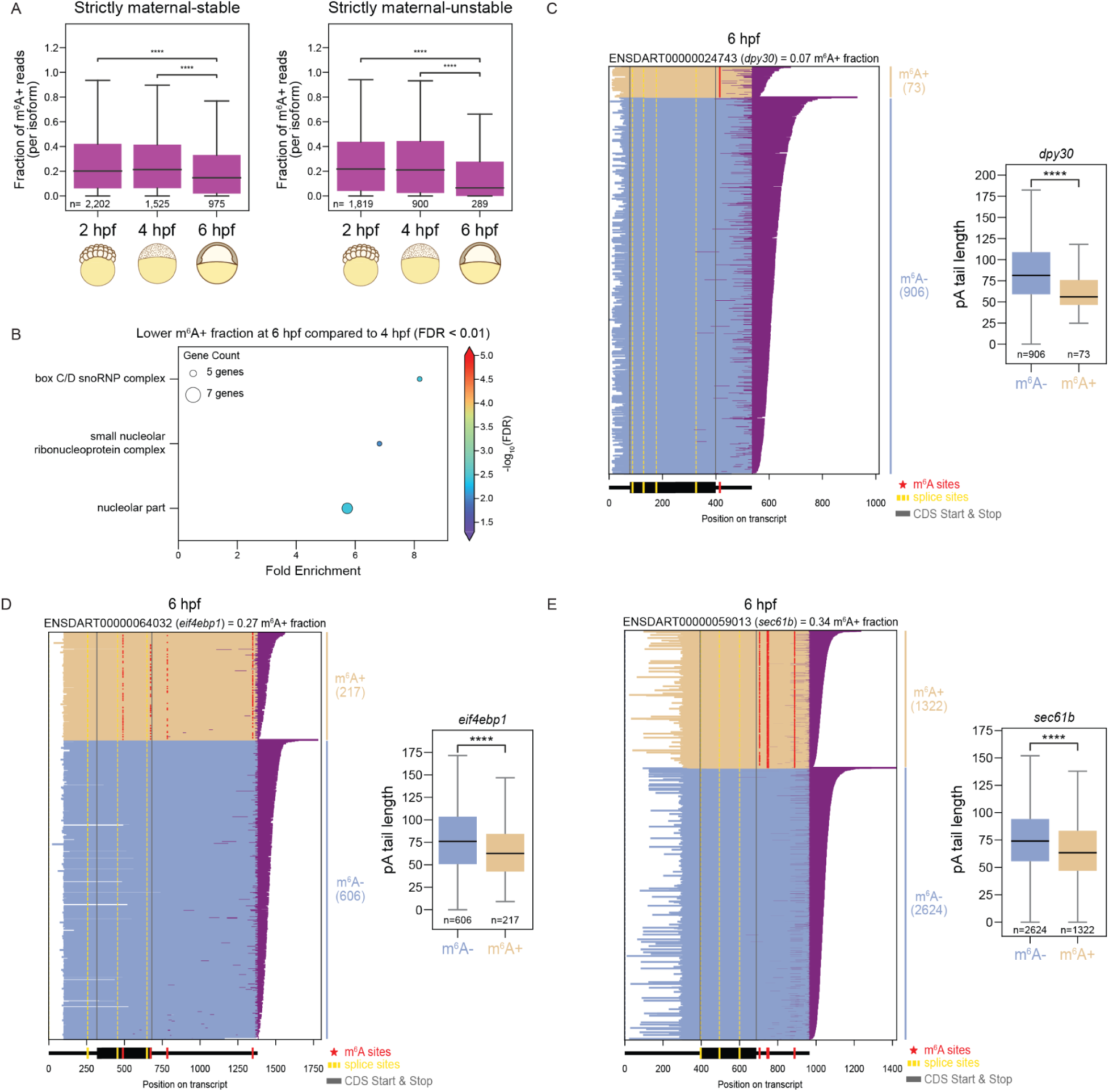
m^6^A+ fraction declines in strictly maternal transcripts and read-level examples of m^6^A-associated poly(A) tail shortening at 6 hpf. (**A**) Fraction of m^6^A+ reads per isoform for strictly maternal-stable (left) and strictly maternal-unstable (right) transcripts, classified using the Bhat *et al.* 2023 SLAM-seq dataset^71^, at 2, 4, and 6 hpf. *n* values are indicated below each box. Significance was assessed by Mann-Whitney test with Bonferroni correction (**** P<0.0001). Only uniquely mapping reads were included, and isoforms were required to have at least 25 reads per replicate. (**B**) Gene ontology fold enrichment for genes with a significant decrease in m^6^A+ read fraction from 4 to 6 hpf (FDR < 0.01). Point size indicates gene count and color indicates −log_10_(FDR). (**C-E**) Read-level views of three representative isoforms at 6 hpf, each with poly(A) tails shown (purple) and accompanying box plots comparing mean poly(A) tail length of m^6^A+ (tan) versus m^6^A- (blue) reads: (C) ENSDART00000024743 (*dpy30*), (D) ENSDART00000064032 (*eif4ebp1*), and (E) ENSDART00000059013 (*sec61b*). In each read-level plot, m^6^A sites (red), splice sites (yellow), and CDS start and stop (gray) are indicated below, with m^6^A+ and m^6^A- read counts shown on the right. P-values for the box plot comparisons were calculated by Mann-Whitney U test (**** P<0.0001).

**Supplementary Figure 5.**
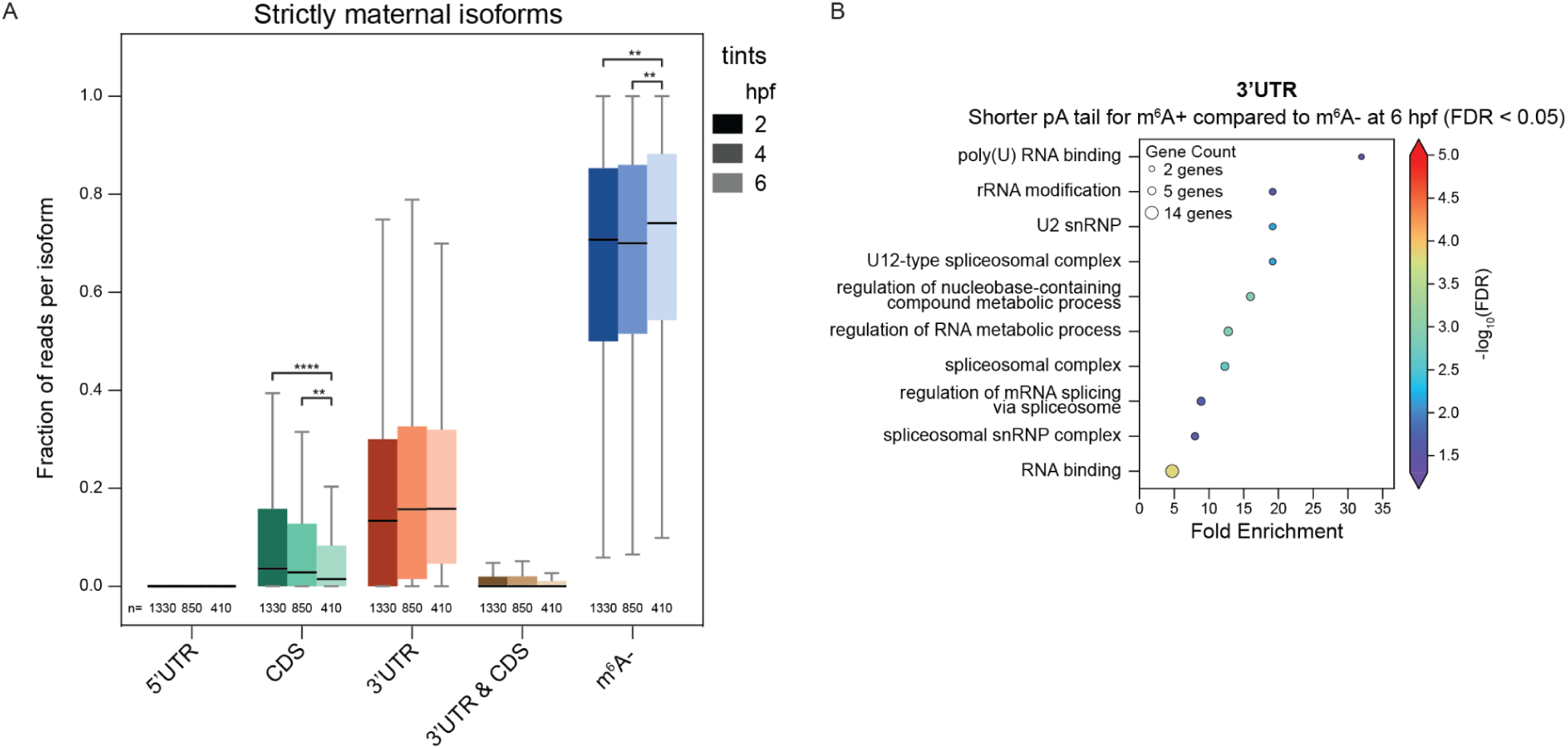
Regional m^6^A decay dynamics are preserved in strictly maternal transcripts. (**A**) Fraction of reads per isoform carrying m^6^A in each transcript region (5′-UTR, CDS, 3′-UTR, or both 3′-UTR and CDS) or lacking m^6^A (m^6^A-), at 2, 4, and 6 hpf (tints), restricted to strictly maternal isoforms defined by Bhat et al. 2023^71^. *n* values are indicated below each box. Statistical significance was assessed by Mann-Whitney U test (** P<0.01; **** P<0.0001). (**B**) Gene ontology fold enrichment for genes with significantly shorter poly(A) tails in m^6^A+ reads relative to m^6^A- reads in the 3′-UTR at 6 hpf (FDR < 0.05). Point size indicates gene count and color indicates −log_10_(FDR).

**Supplementary Figure 6.**
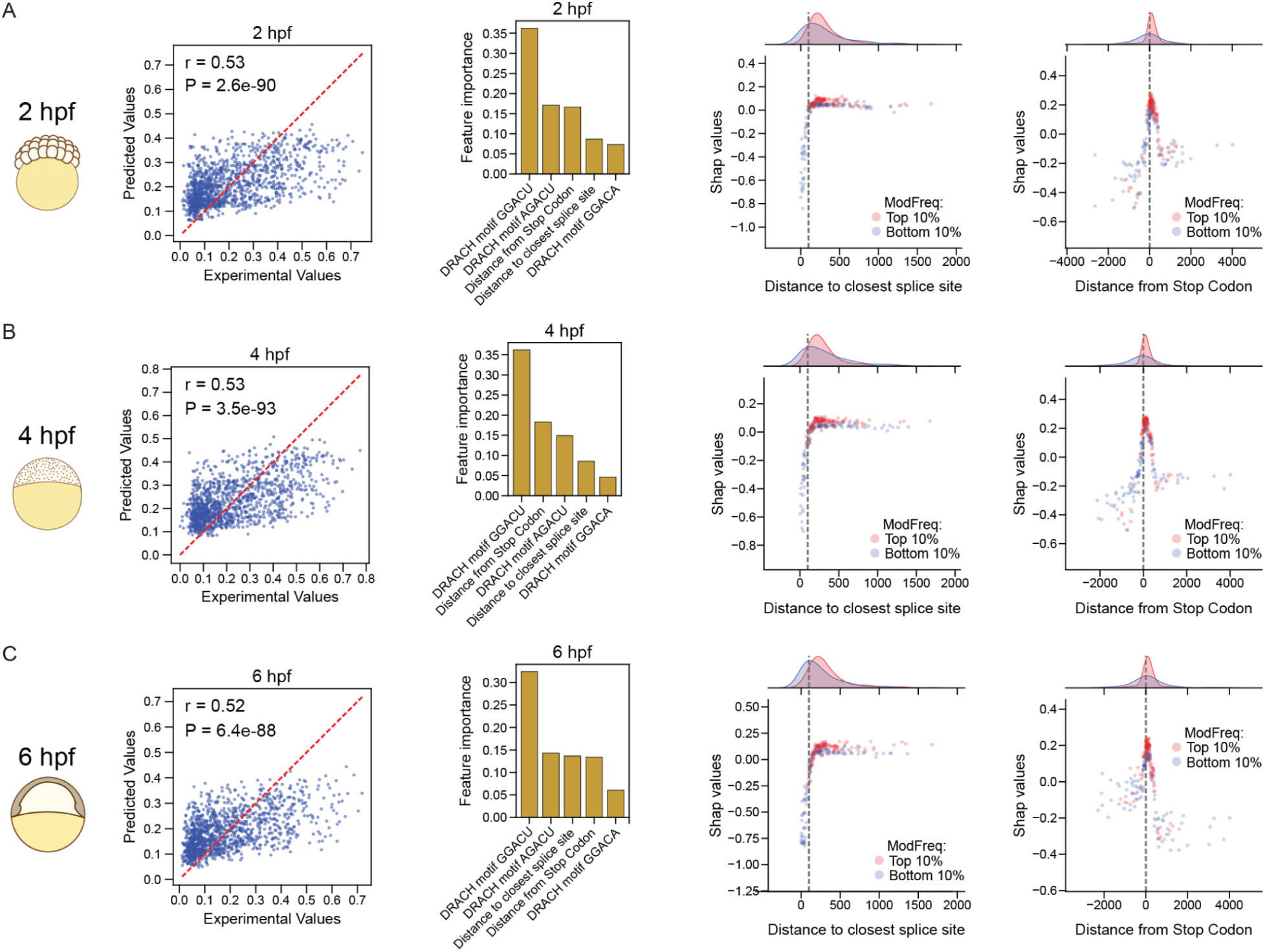
Structural and sequence determinants of m^6^A stoichiometry are stable across MZT. Machine learning prediction of per-site m^6^A stoichiometry from sequence and isoform structure features at 2 hpf (**A**), 4 hpf (**B**), and 6 hpf (**C**). For each time point, from left to right: predicted versus experimental stoichiometry values, with the red dashed line indicating the identity (y = x) line and *r* the Spearman correlation; the top features ranked by importance; SHAP values as a function of distance to the nearest splice site for sites in the top 10% (red) and bottom 10% (blue) of stoichiometry, with marginal density distributions shown above; and SHAP values as a function of distance from the stop codon for the same top and bottom 10% groups, with marginal density distributions shown above.

**Supplementary Figure 7.**
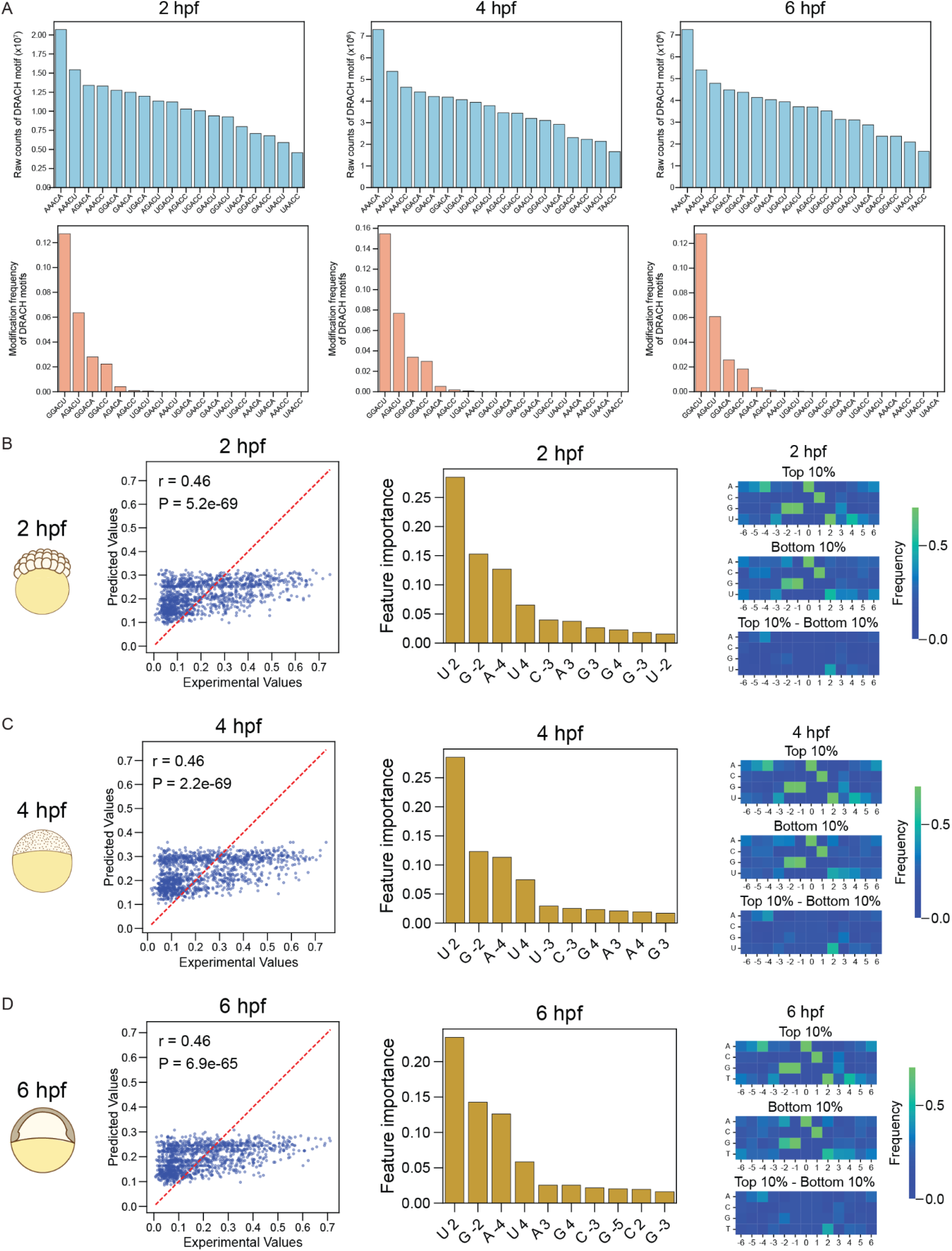
DRACH variant usage and flanking-sequence determinants of m^6^A stoichiometry are consistent across MZT. (**A**) Raw counts of each DRACH motif variant across all isoforms (upper) and the m^6^A modification frequency of each DRACH variant (lower), at 2, 4, and 6 hpf. (**B–D**) Machine learning prediction of per-site m^6^A stoichiometry from the DRACH motif and ±4 flanking nucleotides at 2 hpf (B), 4 hpf (C), and 6 hpf (D). For each time point, from left to right: predicted versus experimental stoichiometry values, with the red dashed line indicating the identity (y = x) line; the top features ranked by importance, labeled by nucleotide and position relative to the methylated adenosine; and nucleotide frequency at each flanking position for sites in the top 10% (upper) and bottom 10% (middle) of stoichiometry, with the frequency difference between the two groups (lower). For the scatter plots, r and the corresponding P-value were derived from Spearman correlation; color in the heatmaps indicates nucleotide frequency.

**Supplementary Figure 8.**
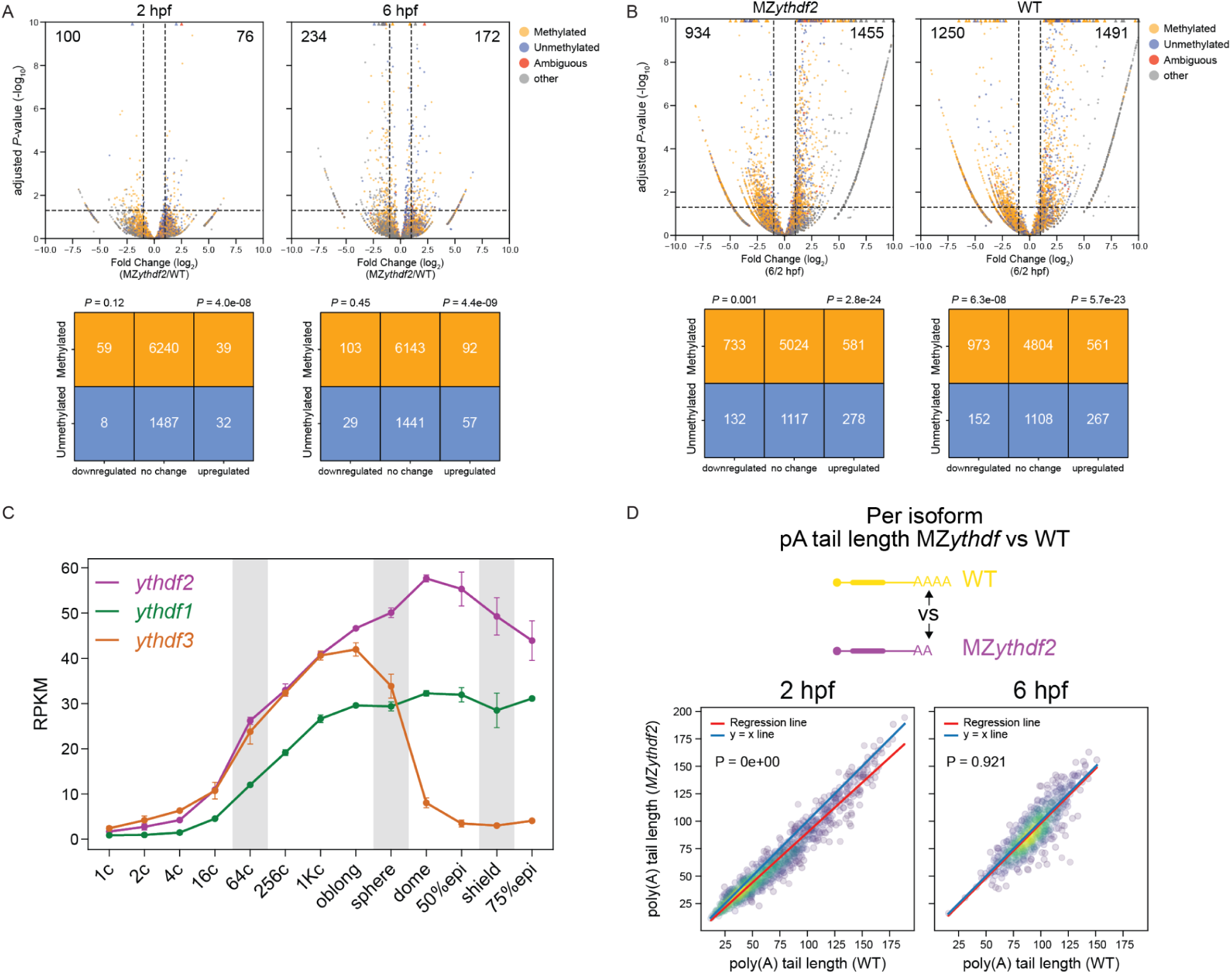
Ythdf paralogs expression across MZT and methylation-resolved differential expression in MZ*ythdf2* and WT sibling embryos. (**A**) Volcano plots of differential gene expression between MZ*ythdf2* and WT sibling embryos at 2 hpf (left) and 6 hpf (right), with genes colored by methylation classification from the WT 2 hpf time course (methylated, yellow; unmethylated, blue; ambiguous, red; other, gray). Points beyond the axis limits are shown as triangles. Lower: contingency tables of methylated and unmethylated genes by differential expression category (downregulated, no change, upregulated). P-values were calculated by chi-square test of proportions, comparing the methylated and unmethylated fractions within each differentially expressed category to those of the remaining genes. (**B**) Volcano plots of differential gene expression between 2 and 6 hpf for MZ*ythdf2* (left) and WT (right), colored and annotated as in (A), with contingency tables and statistics below. (**C**) Expression (RPKM) of the three *ythdf* paralogs (*ythdf2*, purple; *ythdf1*, green; *ythdf3*, orange) across zebrafish development in poly-(A)-selected RNA-seq dataset from Vejnar *et al* ^9^. Stages corresponding to the 2, 4, and 6 hpf DRS time points are shaded in gray. Error bars indicate variation between replicates (standard deviation). (**D**) Per-isoform scatter plots of mean poly(A) tail length in MZ*ythdf2* versus WT at 2 hpf (left) and 6 hpf (right). The red line is the regression fit and the blue line is the identity (y = x) line. Color indicates point density. P-values are shown for the difference between the regression and identity lines (see Methods for calculation details).

